# Characterization of immune cell populations in the tumor microenvironment of colorectal cancer using high definition spatial profiling

**DOI:** 10.1101/2024.06.04.597233

**Authors:** Michelli F. Oliveira, Juan P. Romero, Meii Chung, Stephen Williams, Andrew D. Gottscho, Anushka Gupta, Susan E. Pilipauskas, Syrus Mohabbat, Nandhini Raman, David Sukovich, David Patterson, Visium HD Development Team, Sarah E. B. Taylor

## Abstract

Colorectal cancer (CRC) is the second-deadliest cancer in the world, yet a deeper understanding of spatial patterns of gene expression in the tumor microenvironment (TME) remains elusive. Here, we introduce the Visium HD platform (10x Genomics) and use it to investigate human CRC and normal adjacent mucosal tissues from formalin fixed paraffin embedded (FFPE) samples. The first assay available on Visium HD is a probe-based spatial transcriptomics workflow that was developed to enable whole transcriptome single cell scale analysis. We demonstrate highly refined unsupervised spatial clustering in Visium HD data that aligns with the hallmarks of colon tissue morphology and is notably improved over earlier Visium assays. Using serial sections from the same FFPE blocks we generate a single cell atlas of our samples, then we integrate the data to comprehensively characterize the immune cell types present in the TME, specifically at the tumor periphery. We observed enrichment of two pro-tumor macrophage subpopulations with differential gene expression profiles that were localized within distinct tumor regions. Further characterization of the T cells present in one of the samples revealed a clonal expansion that we were able to localize in the tissue using in situ gene expression analysis. In situ analysis also allowed us to perform in-depth characterization of the microenvironment of the clonally expanded T cell population and we identified a third macrophage subpopulation with gene expression profiles consistent with an anti-tumor response. Our study provides a comprehensive map of the cellular composition of the CRC TME and identifies phenotypically and spatially distinct immune cell populations within it. We show that the single cell-scale resolution afforded by Visium HD and the whole transcriptome nature of the assay allows investigations into cellular function and interaction at the tumor periphery in FFPE tissues, which has not been previously possible.

## Introduction

Colorectal cancer (CRC) accounted for 9.4% of cancer-related deaths (0.9 million) in 2020, and its global incidence is predicted to double by 2035^1,2^. Its poor overall 5-year survival rate highlights the need for better early detection and prognostic biomarkers that can be used in future disease management strategies^3^. During the past decade, there has been growing evidence that tumor heterogeneity is best described at the transcriptome level, rather than with classical histological or mutation-centered disease classifications^4^. Therefore, technologies that refine our understanding of the tumor microenvironment (TME), including the diverse roles of innate and adaptive immune responses and cellular crosstalk in CRC, have the potential to inform better clinical intervention strategies.

Sequencing-based genomics technologies have played an important role in building our current knowledge of CRC biology^4–7^. However, bulk sequencing approaches, which average the data from cells and tissues, are confounded by the complexities of the tumor microenvironment (TME) and intratumor heterogeneity. Single cell transcriptomics (scRNA-seq) technologies have in part filled this gap and allowed for detailed exploration of the cell types within the TME in CRC^8–16^. While these studies add critical single cell level resolution to our understanding of CRC, they lack any information about the organization of the cells within the tissue. Spatial transcriptomics technologies offer a solution. Several commercial technologies are currently available for discovery-based spatial transcriptomics, including Visium CytAssist Spatial Gene Expression (“Visium v2”, 10x Genomics), STOmics (BGI), and Curio Seeker (Curio Bioscience). Other published methods include Seq-Scope^17^, Nova-ST^18^, Open-ST^19^, HDST^20^, DBiT-seq^21^, Pixel-seq^22^, and XYZeq^23^. These methods have enabled the localization of cell types within tissues, which is critical for understanding the interaction between cells in the TME of CRC^24–27^. However, these technologies lack resolution at the single cell scale, or are typically only compatible with fresh frozen tissues, and as such, a deep understanding of tumor organization based on readily available biobanked samples remains elusive.

Here, we introduce Visium HD Spatial Gene Expression (“Visium HD”) and demonstrate its use as a discovery platform for profiling CRC in multiple patients using FFPE tissue blocks. Visium HD slides provide a dramatically increased oligonucleotide barcode density over the Visium v2 slides (11,000,000 continuous features in a 6.5 mm Visium HD capture area, compared to 5,000 features with gaps between them in a 6.5 mm Visium v2 capture area). The single cell scale resolution of Visium HD allowed us to map distinct populations of immune cells, specifically macrophages and T cells, and evaluate differential gene expression at the tumor boundary to explore the potential contribution of these immune cell populations in the TME.

Using an FFPE compatible single cell workflow (the probe-based, Chromium Single Cell Gene Expression Flex) we also generated a multi-patient single cell reference dataset from a larger cohort of FFPE samples and used it to refine our ability to identify distinct cell types. We used this dataset to deconvolve the Visium HD data bins, validating the cell type populations identified by Visium HD and subsequently using the integrated data to comprehensively map the cellular composition and molecular signatures of the TME in CRC. To better understand the interaction between the tumor and its surroundings, we examined the peripheral region that surrounds the tumor by 50 µm. This analysis allowed us to spatially map distinct subpopulations of macrophages to specific regions of the tumor, and compare their transcriptomic profiles which indicate they may exert pro-tumor roles via different pathways. This level of TME characterization was only possible at the resolution of Visium HD, which allowed us to specifically interrogate the cells in closest proximity to the tumor which are likely to have the greatest impact on tumor progression.

With any new technology, validation of findings with an orthogonal approach is critical. To validate the spatial accuracy of Visium HD, we analyzed a subset of genes using an independent spatial technology (Xenium In Situ Gene Expression) and saw strong concordance between the different technology readouts. Next, we mapped macrophages, tumor subpopulations, and T cells that we had observed in the TME via Visium HD data, at single cell resolution using the Xenium technology. Xenium corroborated the presence of the two pro-tumor macrophage subpopulations in different niches and allowed us to pinpoint the location of the T cells within the TME. Using Xenium, we were also able to detect a clonally expanded T cell population and the cellular microenvironment in which it resides, revealing an anti-tumor niche with a third macrophage subpopulation.

Our study underscores the importance of using high resolution spatial technologies in exploring the heterogeneity of tumor biology. These advanced tools are crucial for precisely mapping the diverse immune cell niches within CRC and elucidating the complex interactions between these cells and their microenvironment. By leveraging spatial technologies, we can gain a detailed understanding of spatial variations in cell types and subpopulations, and cell-to-cell relationships, which are key to developing targeted therapies and personalized medicine approaches. The combination of whole transcriptome and targeted in situ spatial technologies used in our investigation provides deeper insights into the complex and dynamic nature of the TME, highlighting the importance of spatial context in understanding cancer heterogeneity and progression.

## Results

### Visium HD specifications and performance

In this study, we included five patients with colorectal adenocarcinoma (Table 1), from which we obtained FFPE (CRC n = 5, and normal adjacent tissue (NAT) n = 3) blocks. Serial sections of FFPE tissues were prepared and selected samples were included to benchmark the technology performance or to explore the TME using Visium HD. Additionally, selected serial sections from the same FFPE blocks were used to generate a single cell RNA-seq dataset and for evaluation via in situ gene expression (Figure 1).

**Figure 1.**
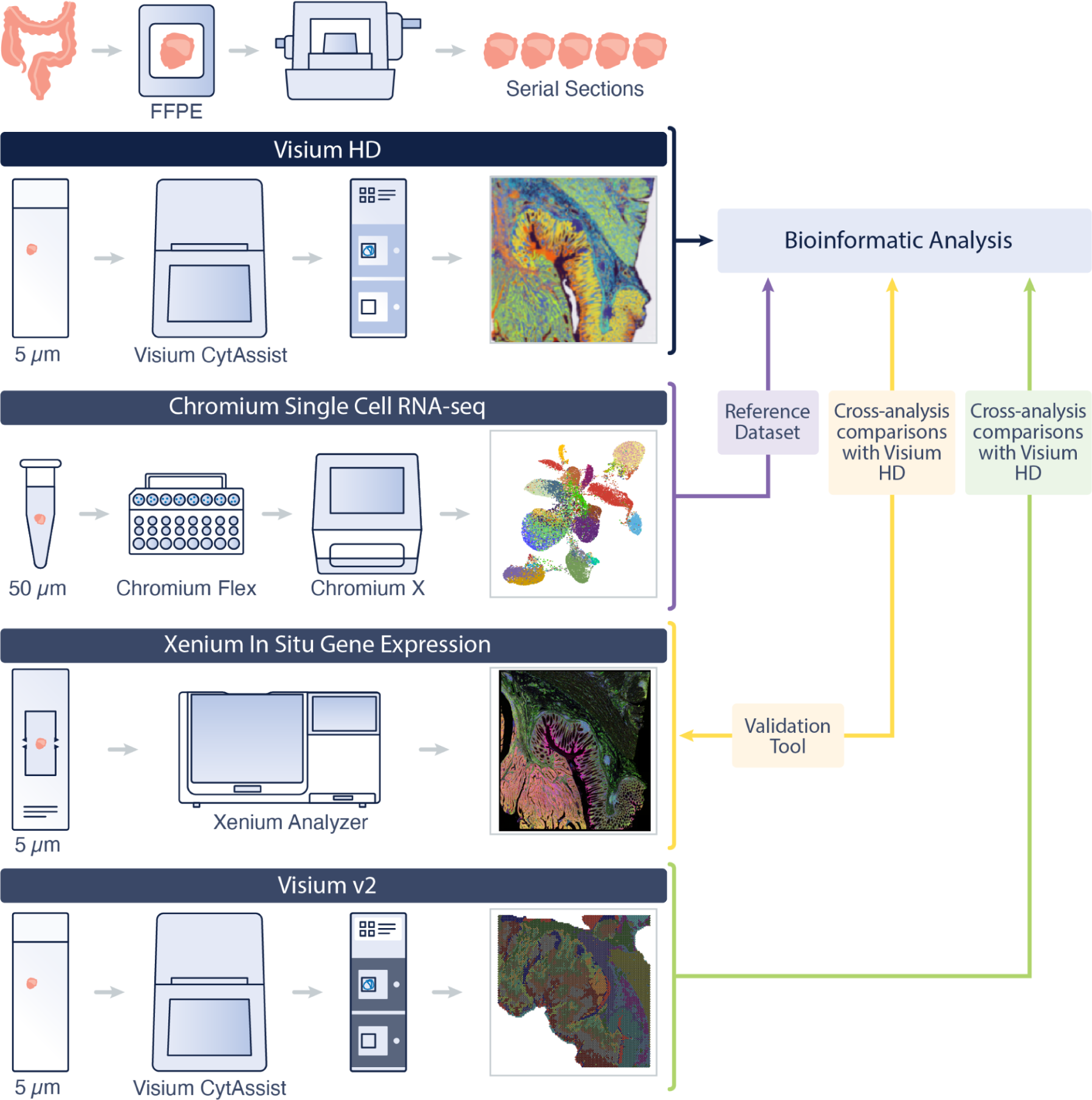
Analysis of CRC and NAT samples using Visium HD. Serial tissue sections were taken from colorectal adenocarcinoma (CRC, n = 5 samples) and normal adjacent tissues (NAT, n = 3 samples) FFPE blocks. A subset of samples were selected and analyzed with the Visium HD assay (n = 3 CRC and n = 2 NAT). Sections from the same FFPE blocks were assayed with single cell RNA-seq (Chromium Single Cell Gene Expression Flex; n = 8). Serial sections were analyzed with Xenium In Situ gene expression (n = 4 CRC) and assayed via the Visium v2 assay (n = 1 CRC and n = 2 NAT). Single cell data were used to create a reference dataset for cell type annotation. In situ data were used for validation of the findings from the Visium HD data and for subsequent analyses. Technology performance comparisons were performed using data from matched datasets.

**Table 1.** Samples evaluated in this study.

| Sample | Colon region | Stage | Sex | Age | Visium HD | Visium v2 | Xenium | Chromium Flex | Chromium TCR |
| --- | --- | --- | --- | --- | --- | --- | --- | --- | --- |
| P1 CRC | Sigmoid | II-A | F | 72 | Included |  | Included | Included |  |
| P2 CRC | Sigmoid | Unknown | M | 60 | Included | Included | Included | Included |  |
| P2 NAT |  |  |  |  |  |  |  | Included |  |
| P3 CRC | Transversum | Unknown | M | 83 |  |  |  | Included |  |
| P3 NAT |  |  |  |  | Included | Included |  | Included |  |
| P4 CRC | Not specified | II-A | F | 61 |  |  |  | Included |  |
| P5 CRC | Not specified | IV-A | F | 58 | Included |  | Included | Included | Included |
| P5 NAT |  |  |  |  |  | Included |  | Included |  |

CRC = Colorectal cancer, NAT = normal adjacent tissue. NAT was available for P2, P3 and P5.

The Visium HD assay enables spatial gene expression analysis with probes targeting the whole transcriptome at single cell scale. Visium HD slides contain two 6.5 x 6.5 mm capture areas within a 8 x 8 mm fiducial frame, where each capture area consists of ∼11 million 2 x 2 µm squares arranged in a continuous array of uniquely barcoded oligonucleotides (Figure 2A). Importantly, the 2 µm squares are directly adjacent to each other, resulting in a continuous lawn of capture oligonucleotides with no gaps between features, representing an improvement over earlier Visium slides, which have 55 µm circular capture areas with gaps between them (Figure 2B). For downstream analysis, the 2 µm data can be used directly or collated into larger bins to increase the coverage of the data; the Space Ranger (v3.0) pipeline outputs the 2 µm data and data binned at 8 and 16 µm resolution (unless otherwise described, the 8 µm binned data were used in this study). To assess the increased resolution afforded by Visium HD, we analyzed serial sections from a normal colon mucosa sample run on Visium v2 and Visium HD. Visium HD generated notably higher resolution data, as shown in the improved unsupervised clustering, both in terms of the total number of clusters detected (18 clusters in Visium HD vs. 3 clusters in Visium v2), and the ability to map them to morphological features of the colon mucosal tissue (Figure 2C). Next, we assessed the correlation between Visium v2 and Visium HD data using serial sections of colon samples (2 NAT samples and 1 CRC sample, Figure 2D and Supplemental Figure 1). Our data show a strong correlation between UMI counts at the whole transcriptome level at similar sequencing depth across an entire tissue (matched tissue areas), highlighting that the data obtained from each assay are highly comparable in terms of sensitivity across the colon tissues analyzed (R^2^ = 0.82, Figure 2D; R^2^ = 0.81 and 0.90, Supplemental Figure 1). To remove any potential bias arising from off-target probe binding events, i.e., probes binding to genomic DNA (gDNA), which could introduce sensitivity biases in this analysis, we compared the UMI counts for the subset of probes that spanned only exon-exon junctions (7,605 probes out of 54,580). The estimated fraction of molecules (measured by the number of UMIs) arising from gDNA was 4.1% in Visium v2 and 0.5% in Visium HD. When comparing counts from the probes spanning exon-exon junctions, we observed a stronger correlation between assays (R^2^ = 0.92, Figure 2D; R^2^ = 0.93 and 0.96, Supplemental Figure 1), indicating that the increased resolution of the Visium HD assay maintains the high assay sensitivity of Visium v2.

**Figure 2.**
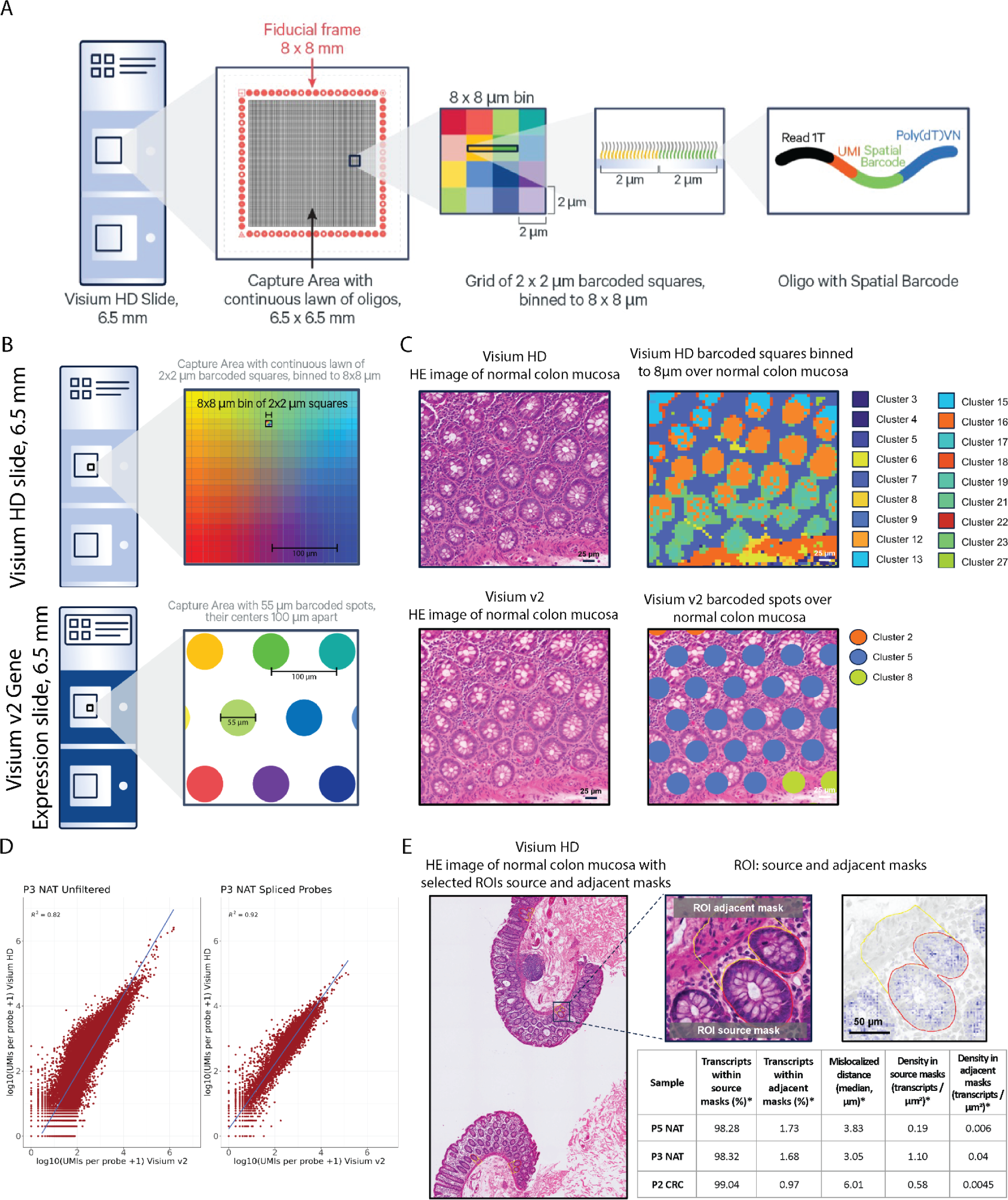
Visium HD Spatial Gene Expression slide architecture and performance. **A.** Visium HD slide with two 6.5 x 6.5 mm capture areas, each containing a continuous lawn of uniquely barcoded 2 x 2 µm squares, which are binned to 8 µm squares for downstream analysis. **B.** Visium HD slides, compared to Visium v2, which have spots of 55 µm diameter spaced 100 µm apart. **C.** Comparison of serial sections of a representative normal colon mucosa sample P3 NAT. Visium HD detects eighteen clusters that closely correspond to tissue morphology, while Visium v2 detects three clusters. **D.** Sensitivity comparison between Visium HD and Visium v2 on representative sample P3 NAT. Left plot shows expression levels of all probes (whole transcriptome); the right plot shows only probes spanning an exon-exon splice junction. Diagonal lines represent x = y. **E.** Transcript localization accuracy analysis performed across four randomly selected regions of interest (ROIs) per tissue section for selected goblet cell gene markers (*CLCA1, FCGBP, MUC2*); source masks are colon gland structures, adjacent masks are the immediately adjacent regions containing lamina propria. Images show selected ROIs in a representative normal sample P3 NAT; red lines outline the source mask, yellow lines the adjacent mask. Table shows the median percentage of localized transcripts in the source and adjacent masks, the density of selected transcripts in both masks, and the distance of selected transcripts from source masks (*). Four ROIs in each colon sample were included in this analysis.

In array-based spatial technologies, mRNA must migrate from the tissue to come into contact with a primer (or *vice versa*, the primer must come into contact with the mRNA molecule). However, because transcript migration does not occur linearly, if the tissue placement and subsequent molecular biology reactions are not carefully controlled, the spatial accuracy of mRNA detection may be impacted, i.e., transcripts may be detected away from their origin. Similar to the Visium v2 assay, Visium HD utilizes a controlled environment to transfer analytes from tissues to the capture arrays (the CytAssist instrument), improving spatial accuracy of RNA detection^28,29^. Poor transcript spatial accuracy has the potential to impact biological interpretations, thus we sought to assess this in samples run through the Visium HD workflow. For this analysis, we evaluated two NAT samples and one CRC sample (Figure 2E). We selected genes that are known to be localized within glands of normal colon mucosal tissue (goblet cell gene markers: *CLCA1, FCGBP, MUC2*). In each tissue section, we manually selected four random regions of interest (ROIs) matching colon glands (“source masks”) and their immediate adjacent regions containing lamina propria (“adjacent masks”) and measured the transcript localization accuracy of the selected goblet cell gene markers. Across each sample, the majority of transcripts were localized in their expected morphological locations within the source masks (98.3 – 99%), and only a small proportion (0.97 – 1.73%) were in adjacent masks (Figure 2E), demonstrating the high spatial accuracy of mRNA detection obtained from Visium HD.

### Visium HD reveals the spatial landscape of CRC tumors at single cell scale

To characterize the spatial landscape of the CRC samples, sections from three blocks (P1 CRC, P2 CRC and P5 CRC) were selected for profiling using Visium HD. The resulting unsupervised clusters aligned with the expected morphological features, highlighting the spatial organization of the samples (Figure 3A). Since each section was analyzed independently, the results were patient specific and thus limited our ability to perform cell type comparisons between sections. To ensure we had the most refined and consistent cell type labels across all samples, we generated a single cell reference atlas from serial FFPE sections of CRC and NAT (n = 8 blocks, Table 1), which included the same three blocks selected for Visium HD. This approach enabled us to sample and profile 245,494 cells (after quality control analysis), which gave us more power for cellular annotations. We then applied differential gene expression (DGE) analysis to identify marker genes between these clusters. We manually classified the graph-based clusters into ten broad cell types, denoted as level 1 annotations. For level 2 annotations, we repeated the clustering process within each level 1 cluster, maintaining 25 PCs but adjusting the resolution parameter to 0.1 to prevent over-splitting. We identified marker genes through DGE and annotated cell types manually based on published cell gene markers (Supplemental Figure 2). We then used this annotated single cell dataset as a reference to deconvolve the HD data, assigning a homogeneous set of cell type labels across the different samples (See Methods and Figure 3B). To compare the Visium HD unsupervised clustering results with the deconvolved labels, we plotted confusion matrices for each sample as heatmaps and observed that the most prominent cell types were also detected by unsupervised clustering (Figure 3C). These findings confirm that the expected cell types in the colon mucosa can be identified based on the Visium HD data alone and deconvolution using single cell data is not required. However, deconvolution based on single cell data and assignment of uniform labels is useful for performing comparisons between samples.

**Figure 3.**
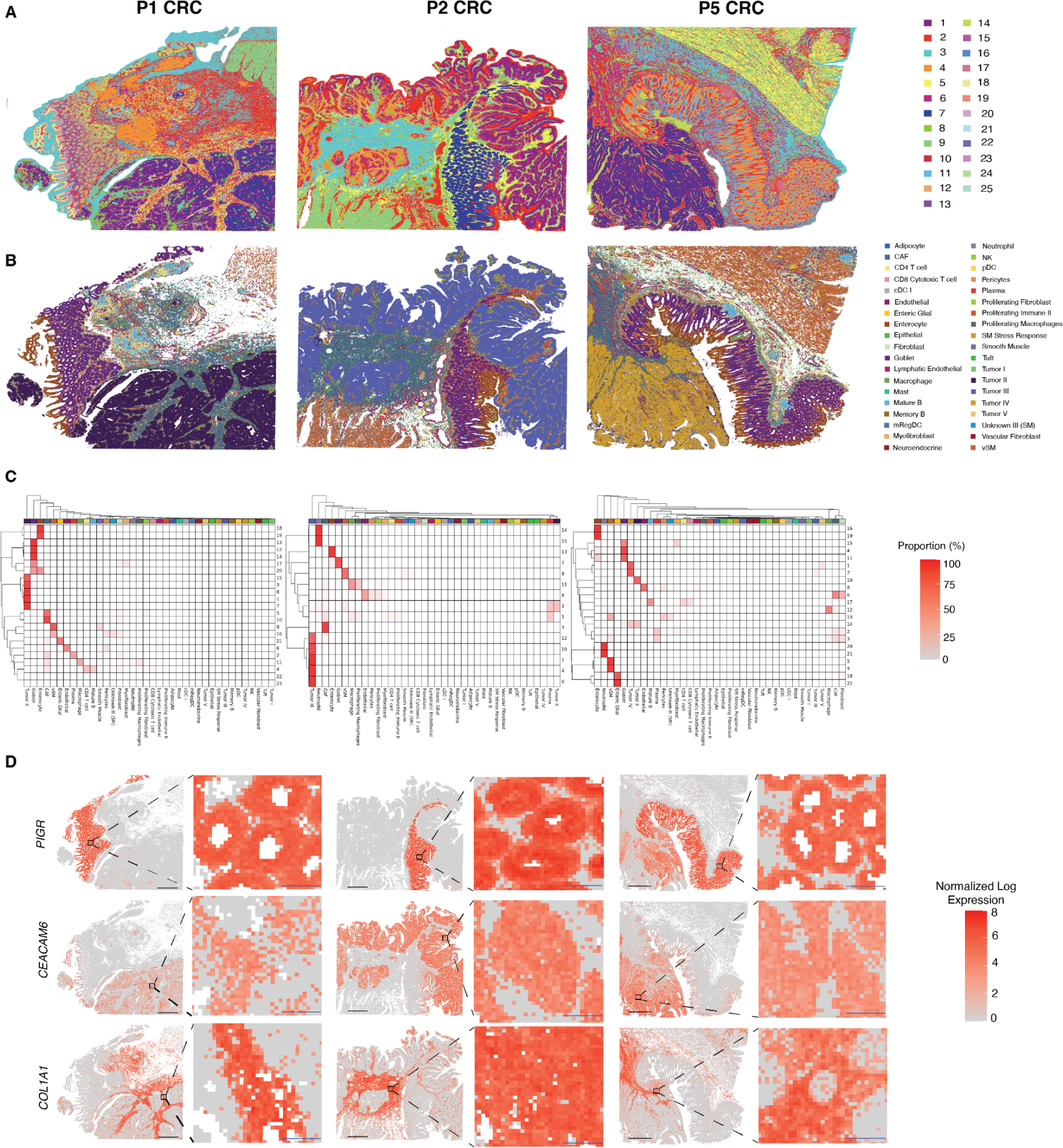
Spatial mapping of CRC samples using Visium HD reveals high resolution, accurate transcript mapping. **A.** Spatial mapping of three CRC samples (P1 CRC, P2 CRC and P5 CRC) with 8 µm bins colored based on unsupervised clustering. **B.** Spatial mapping of the same three CRC samples with 8 µm bins colored by cell types predicted by deconvolution using the single cell reference dataset. **C.** Confusion matrices denoting the relationship between the unsupervised clusters (rows) and labels assigned by deconvolution labels (columns). Data is scaled by row. **D.** Validation of selected cellular gene markers with known spatial localization: *PIGR* (goblet cells and enterocytes), *CEACAM6* (tumor) and *COL1A1* (fibroblasts). Samples correspond to those in **A**. For each sample, the tissue-level view is shown on the left, with the inset as a black box, and the inset view is shown on the right. Scale bars: black = 1 mm; blue = 80 µm.

The deconvolved Visium HD data provided a highly resolved map of the cell types observed in the single cell reference data, aligning with tissue morphology. For example, we mapped most goblet cells and enterocytes in the normal mucosa, cancer-associated fibroblasts and tumor cells were mapped to the tumor area, and multiple immune cell types were mapped throughout the tissue sections (Figure 3B). We observed that each patient sample was associated with a major and distinct tumor cell type (Supplemental Figure 3 and Supplemental Figure 4) mapped onto the morphological tumor regions across each tissue section. We validated the spatial arrangement of these cell labels in Visium HD with the expression of well known markers such as *PIGR* (goblet cells and enterocytes)*, CEACAM6* (tumor) and *COL1A1* (fibroblasts) (Figure 3D).

### Macrophages are enriched at the tumor boundary

Given that immune cell dynamics are known to play a key role in CRC progression, we wanted to characterize the immune cell populations within the TME of our CRC samples. We focused on the tumor boundary region so that we could understand immune cell dynamics and function in these tumors. Taking advantage of the improved resolution afforded by Visium HD, we used distance-based analysis to resolve the cellular composition of tumor boundary, an analysis that is not possible to do at the resolution of Visium v2. We selected all barcoded 8 µm bins within 50 µm of the regions we had labeled as tumor cells via spot deconvolution (Figure 4A) that include only a determined single cell type (i.e. not a mixture of, or undetermined cell types). Once the set of barcodes in these tumor peripheral regions was defined, we quantified the composition of cell types present in the 50 µm region peripheral to the tumor. When compared to the rest of the tissue, cancer associated fibroblasts (CAF) were the most prominent cell type, while macrophages were consistently identified as the most abundant immune cell type, across all tissue blocks studied (Figure 4B). We corroborated these findings, which were derived from the cell type annotation, by examining the expression of known macrophage (*C1QC*) and CAF (*COL1A1)* markers (Figure 4A).

**Figure 4.**
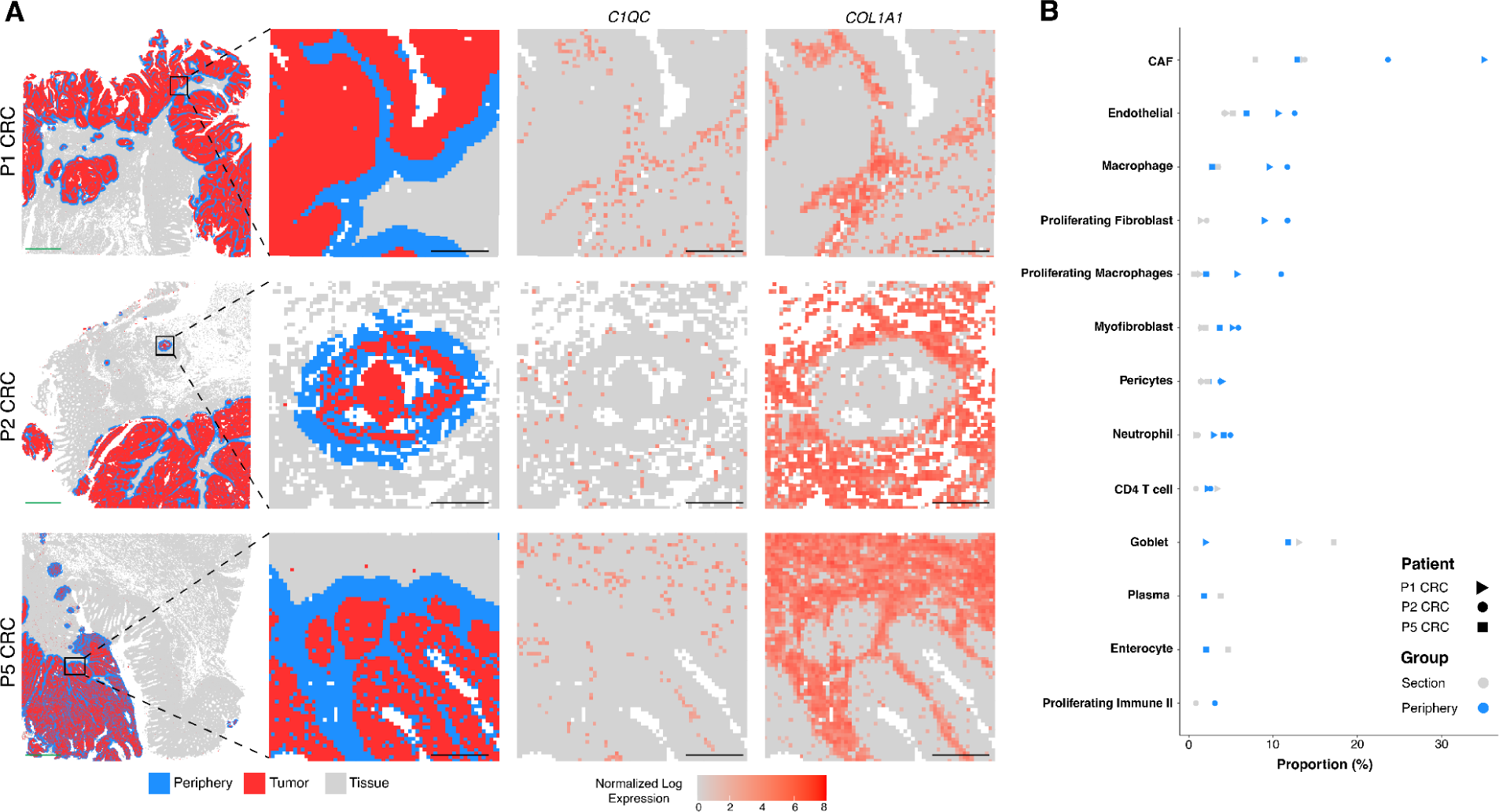
Cellular composition of the tumor periphery in each CRC section. **A.** Analysis of the tumor periphery. 8 μm bins annotated as tumor cells are shown in red, with bins within 50 µm of the tumor periphery shown in blue. Rows correspond to three different samples. The first column shows the 6.5 x 6.5 mm capture area, the second column shows the zoomed in view, the third column shows the corresponding expression of *C1QC* (macrophages), and the fourth column shows the corresponding expression of *COL1A1* (fibroblasts). Scale bars: green = 1 mm; black = 125 µm. **B.** Dot plot with the proportion of cell types in the tumor periphery (blue) and the rest of the tissue section (gray) for the three different blocks.

### Transcriptomic analysis of the macrophage-enriched tumor regions reveals two macrophage subpopulations

As the most abundant immune cell type in the tumor periphery, we focused our analysis on the tumor regions enriched with macrophages to gain insights on their interplay with the TME. First, we evaluated if these cells presented heterogeneous gene expression signatures and spatial locations within the tumor region. To do this, we selected the 8 µm bins deconvolved as macrophages around the tumor region to perform an independent unsupervised clustering analysis. We found two macrophage subpopulations with specific gene expression profiles mainly defined by expression of *SELENOP* or *SPP1* genes (Figure 5A). We then took advantage of the whole transcriptome nature of the assay and performed an enrichment analysis of the differentially expressed genes to further characterize these macrophage subpopulations. We observed that *SELENOP^+^*macrophages were differentially enriched for pathways such as TNF-α signaling via NFK-β, apoptosis pathways, and UV response to DNA damage. Meanwhile, *SPP1^+^* macrophages were enriched for coagulation, cholesterol homeostasis, and upregulation of KRAS signaling pathways (Figure 5B).

**Figure 5.**
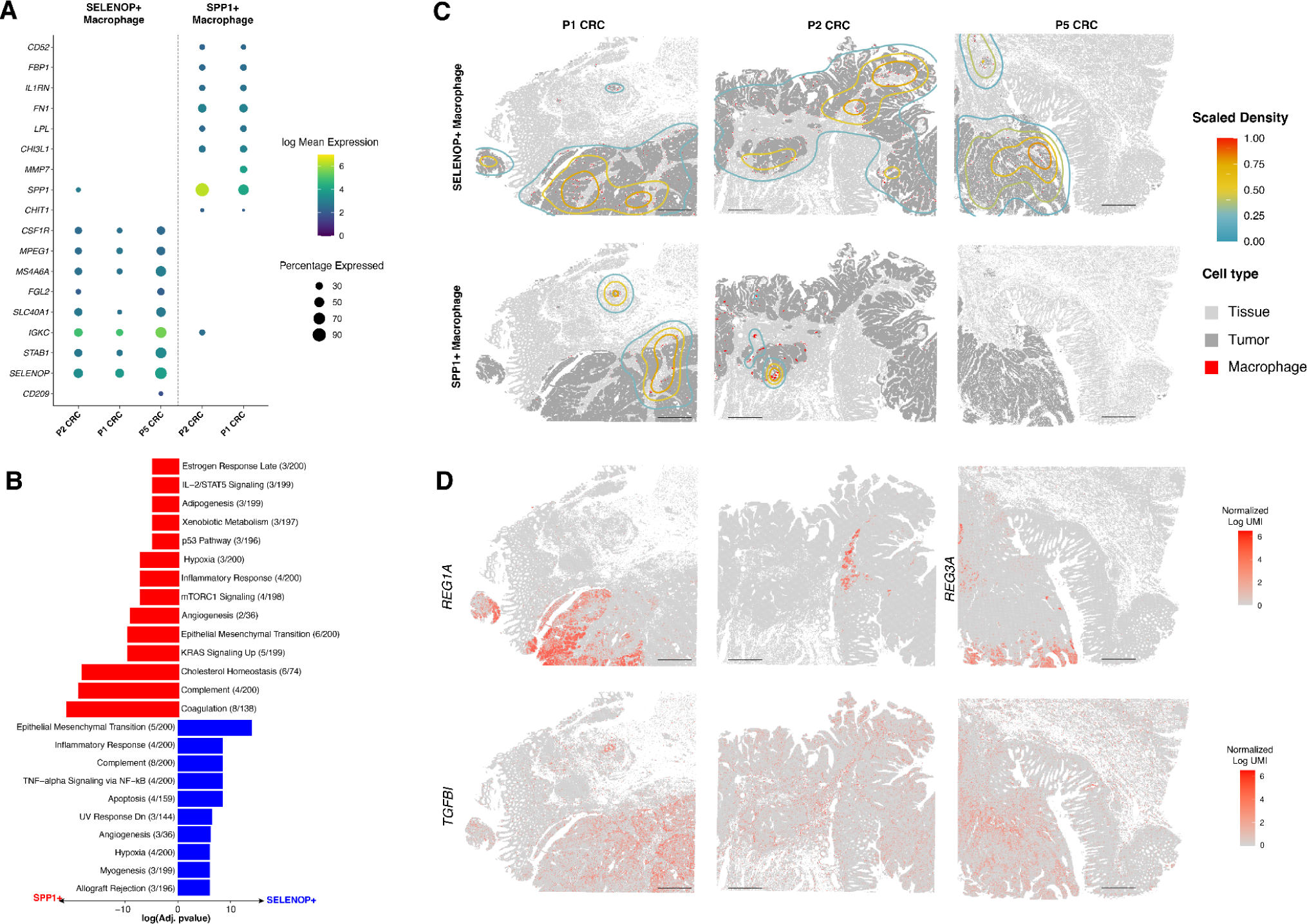
Identification and localization of two macrophage subpopulations in the tumor microenvironment. **A.** Dot plot showing expression profiles of two distinct macrophage subpopulations identified at the boundary in the tumor samples studied. **B.** Bar plot showing the enriched gene sets for the macrophage subpopulations identified. **C.** Kernel density maps showing the differential spatial localization of *SELENOP^+^* and *SPP1^+^* macrophages and how they are associated with tumor areas. **D.** Heatmap showing the expression of REG family genes (*REG1A or REG3A)* and *TGFBI* in the CRC sections. Scale bar = 1 mm.

To add the spatial context of these macrophage subpopulations in the TME, we identified highly enriched regions using density estimation (see Methods) and observed that the *SELENOP*^+^ and *SPP1*^+^ macrophages were mostly in different spatial niches in the tissue (Figure 5C). Analysis of the gene expression profiles of the tumor cells close to these macrophage subpopulations revealed that the different macrophage subpopulations were localized in tumor regions with differential gene expression profiles. Tumor cells in areas enriched for *SPP1^+^* macrophages showed differential expression of *TGFBI*, while tumor regions closer to *SELENOP^+^* macrophages were enriched for *REG1A* and *REG1B* (Figure 5D). Both *TGFBI* and the *REG* gene families have been implicated in tumor progression.

### Characterization and spatial localization of T cells in the TME

The recruitment and function of T cells into the TME has been suggested to be associated with the dynamics of the cells in the tumor niches, and has long been associated with favorable disease outcomes^30^. As with most solid tumor types, CRC tumors are typically cold tumors, which have implications to immunotherapy interventions^31^. However, since our investigation employs high resolution spatial technologies, we wanted to leverage this to specifically explore T cell localization and behavior at the tumor boundary. In our initial analysis, we only included bins predicted to contain only one cell type (singlet 8 µm bins). However, we observed that the tumor periphery region (50 µm around the tumor) displayed enrichment in the number of bins labeled as doublets (two cell types co-existing in the same bin) compared to the rest of the tissue (Figure 6A). This finding is expected, given the known cellular heterogeneity at the boundary of morphologically distinct regions. We found that most T cells were assigned to doublet bins or rejected (the algorithm was unable to predict the cell type), and therefore excluded from our initial analysis, making it more challenging to spatially localize these immune subpopulations (Supplemental Figure 5). To overcome this, we first identified regions enriched in either CD4^+^ or CD8^+^ T cells, independent of whether they were assigned to a singlet or doublet bin (Figure 6B), and performed nuclei segmentation on these regions. We then leveraged the higher resolution 2 µm binned data and assigned the corresponding 2 µm bins that were located within the nuclei polygons to create a gene by nuclei UMI count matrix for further processing. Following this strategy, we were able to identify T cells at the tumor boundary (Figure 6C), but observed that cells expressing *CD8A* and *CD4* were sparsely distributed. We also examined the expression of known T cell markers in this region (*TRAC, CD3*) and other cell type markers such as *PECAM1* (endothelial), *IGKC* (plasma), *COL1A1* (CAF), *SPP1* or *SELENOP* (macrophages), and *CEACAM5* (tumor) (Supplemental Figure 6) to obtain a fine grained map of the cell types in these areas of the tissue. This analysis allowed us to identify and localize both CD4 and CD8 T cells at the tumor periphery but not in the surrounding normal tissue, suggesting that these infiltrating lymphocytes may be playing an active anti-tumor role.

**Figure 6.**
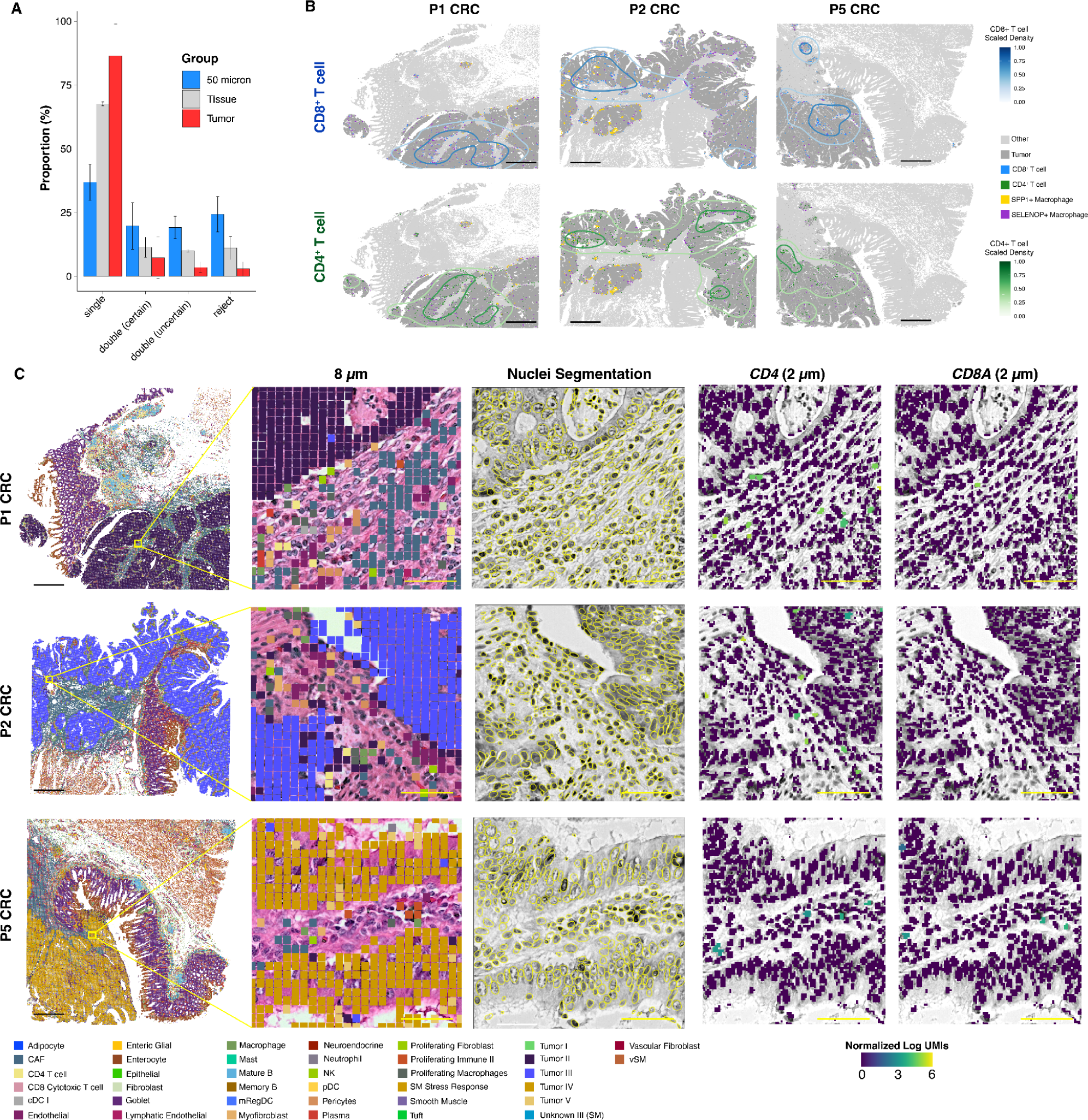
Spatial localization of T cells in the tumor microenvironment. **A.** Barplot showing the proportion of each 8 µm bin class (singlet, doublet, rejected) for each tissue region. **B.** Density maps showing the spatial location of CD8+ and CD4+ T cells in the different samples. **C.** Zoomed-in view of regions with bins labeled by deconvolution results at 8 µm (left), nuclei segmentation results in the zoomed-in regions (center) and normalized expression of *CD4* and *CD8A* (right) of the transformed UMI matrix by grouping 2 µm bins within each of the segmented nuclei. Scale bars: black = 1 mm; yellow = 50 µm.

### Xenium in situ analysis validates Visium HD findings and reveals the spatial distribution of clonally expanded T cells in the TME

To further investigate the spatial distribution of immune cells within the TME and to validate findings from the Visium HD data, we profiled the samples using the Xenium Analyzer. Xenium is an in situ spatial analysis platform that provides subcellular resolution for a targeted set of genes. We have previously shown that Xenium is ∼8.4x more sensitive on a per-gene basis than Visium v2 on a cohort of breast cancer samples^32^, and thus we wanted to use Xenium to validate our Visium HD findings and interrogate the T cell populations more closely. We first set out to benchmark the sensitivity of Xenium with Visium HD in this study. We used the Xenium Human Colon gene expression panel (322 genes) and a custom add-on panel targeting 100 additional genes, which was designed to target diverse immune populations we observed in the Visium HD data (Supplemental Table 1). The panel was used with the Multimodal Cell Segmentation workflow, which allows segmentation of cells based on boundary stains and morphology rather than relying on nuclear expansion alone. To compare the Xenium data to the Visium HD data, we limited the Visium HD data to the 422 genes on the Xenium panel, and found that Xenium was ∼5.7x more sensitive on a per-gene basis. However, when we compare total transcripts identified in the shared region, we see that Visium HD captures ∼6.5x more transcripts than Xenium due to its whole transcriptome nature (Supplemental Figure 7). Both of these results are in line with our previously published comparisons. We expect some differences in sensitivity gains due to differences in the specific genes that are included on the Xenium panels and the nature of the samples used in each study.

We then sought to validate our findings to confirm that the subtypes and localization of macrophages we had observed in the Visium HD data were correct. Consistent with the Visium HD findings, Xenium revealed heterogeneity within both tumor cells and macrophage populations (Figure 7A, 7B). *SELENOP^+^/STAB1*^+^-macrophages were found near *REG1A*^+^ tumor cells (Figure 7C, 7D) while *SPP1^+^* macrophages were localized in close proximity to *TGFBI*^+^-tumor cells (Figure 7E, 7F). Interestingly, we observed that cancer associated fibroblasts (CAFs) which localized at the border of *TGFBI*^+^ tumor also expressed *MMP11* (Figure 7E, 7F), a matrix metalloproteinase that breaks down ECM and is associated with poorer prognosis^33^. This colocalization of *SPP1*^+^ macrophages, *TGFBI*^+^ tumor cells, and *MMP11*^+^ CAFs within the TME may suggest a coordinated effort to promote tumorigenesis.

**Figure 7.**
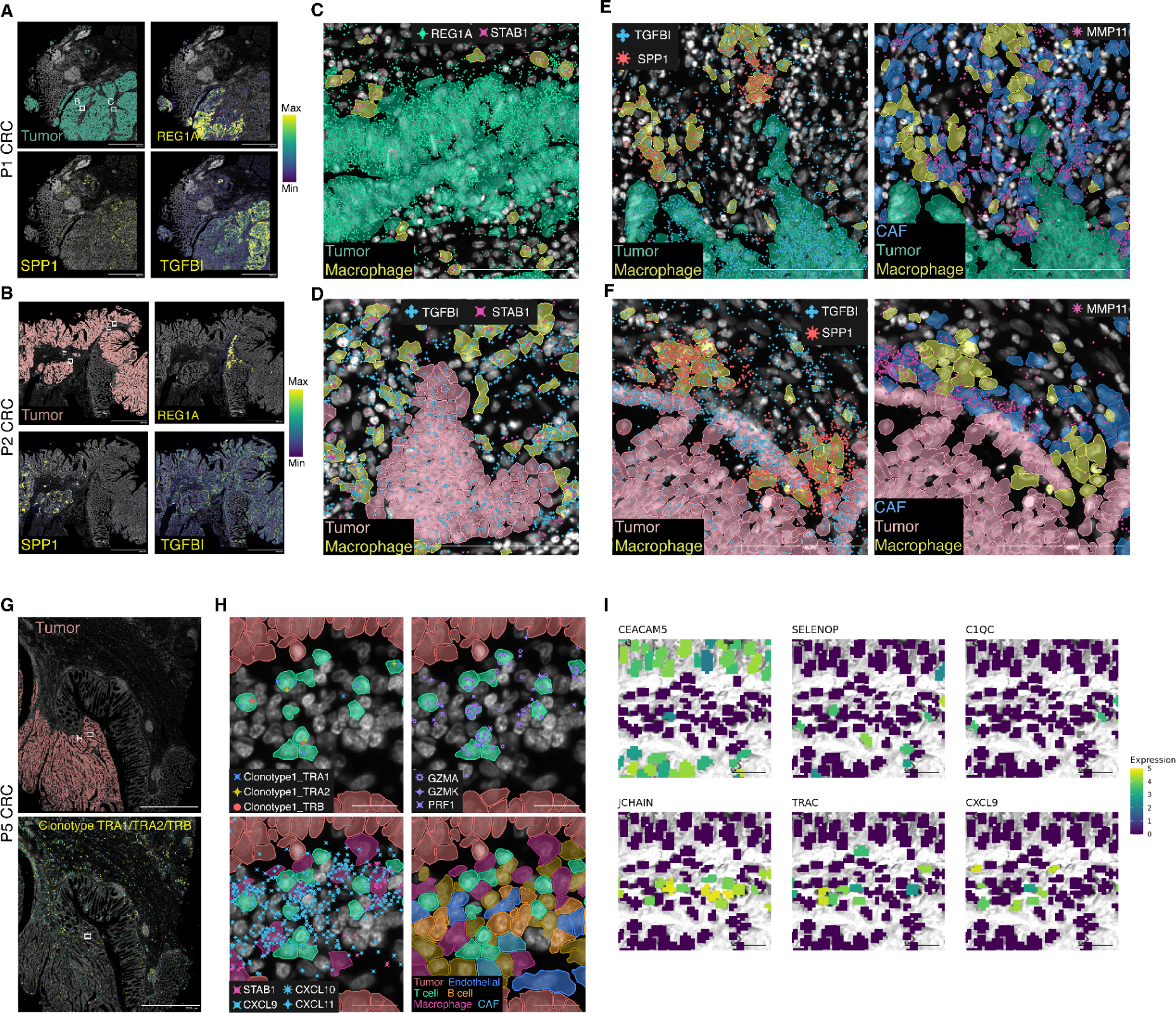
Xenium in situ confirms the existence and localization of macrophage subtypes and clonally expanded T cells in the tumor microenvironment. **A, B.** Expression of *REG1A* and *TGFBI* transcripts (right panels) and *SPP1* (bottom left panel) within tumor region (top left panel). **C, D.** *STAB1*^+^ macrophages near *REG1A*^+^ tumor cells. *STAB1* was used to visualize the macrophage subtype co-expressing *SELENOP*. **E, F.** *SPP1^+^* macrophages shown in proximity of *TGFBI^+^* tumor cells and *MMP11*^+^ cancer associated fibroblasts. **G.** Combined expression of clonotype TRA1/TRA2/TRB in sample P5 CRC. **H.** Clonally expanded CD8 cytotoxic T cells reside closely to tumor cells and within *CXCL9*/*CXCL10*/*CXCL11* foci. **I.** Zoom in view of the same regions using Visium HD with 2 µm bins assigned to segmented nuclei. Bins are colored by the normalized log UMI counts of *CEACAM5*, *SELENOP*, *C1QC*, *JCHAIN*, *TRAC,* and *CXCL9.* Scale bars: 2 mm in A, B, G; 100 µm in C, D, E, F; 20 µm in H; 50 µm in I.

To better understand the T cell response at play we wanted to explore the clonality of the antigen recognizing T cell receptors (TCRs) of the T cells in and around the CRC tumors. To do this, we obtained dissociated tumor cells from the same patient samples and isolated the T cells. We then profiled the TCR clonality of these T cells using the Single Cell Immune Profiling v2 workflow. This analysis revealed a clonotype with 11% representation within the T cell population of sample P5 CRC (TRAV38-1 TRAJ58; TRAB38-2/DV8 TRAJ57; TRBV4-2, TRBJ2-1. Supplemental Table 2), but no expansions in the other samples. To confirm this was a novel clonotype specific to this patient tumor and not present due to on ongoing or prior infection, we searched VDJdb (https://vdjdb.cdr3.net/) and found no known matches to the CDR3 sequences, indicating that this clonotype recognises to a neoepitope specific to this tumor.

As expanded clonotypes demonstrate an active adaptive immune response, we sought to localize these cells within the tissue to better understand the role they were playing in the TME. We designed probes targeting the CDR3 regions of the two alpha and one beta chains of the overlapping expanded clonotypes and included them in the Xenium custom add-on panel (for probe sequences, see Methods). Xenium analysis showed clusters of clonally expanded T cells residing closely to tumor cells and within gut-associated lymphoid tissues (Figure 7G). Gene expression signatures identified these T cells as *CD8*^+^ cytotoxic T lymphocytes (expressing *CD8A*, *PRF1*, *NKG7*, *GZMA*, and *GZMK* genes) (Figure 7H). Interestingly, these T cells were localized within *CXCL9*/*CXCL10*/*CXCL11* foci, where *STAB1*^+^ macrophages, B cells, and endothelial cells are present and contributing to the expression of these chemokines (Figure 7H), known to recruit immune cells to the tumor site^34^. This observation was validated in the corresponding region of the Visium HD data (Figure 7I). *TRAC*^+^ T cells were identified near *CEACAM5*^+^ tumor cells, *SELENOP*^+^/*C1QC*^+^ macrophages, and *JCHAIN*^+^ B cells. *SELENOP* and *JCHAIN* were not included in the Xenium gene panel but we could include them in our analysis based on the Visium HD data, highlighting the complementary strengths of Xenium and Visium HD technologies.

## Discussion

The advent of spatial transcriptomics has enabled a more comprehensive understanding of cellular tissue dynamics in health and disease, and is particularly relevant in the oncology field where the localization of specific cell types in the TME can have prognostic implications. By enabling precise mapping of tumor microenvironments, these technologies reveal the complex spatial relationships and interactions among cells, which are crucial for understanding tumor progression and resistance to therapy. However, existing technologies have limitations related to resolution, tissue compatibility, or ease of use. In this study, we introduced Visium HD, the next generation of the Visium technology, and used it to explore the TME in FFPE colon adenocarcinoma samples.

It has been recognized that the two most important quality parameters of sequencing-based spatial technologies are the sensitivity of mRNA capture per unit area and the spatial accuracy of the mRNA detection^28^. By using a subset of our sample cohort (two NAT and one CRC sample), we demonstrated that Visium HD retained similar gene detection sensitivity when compared to the Visium v2 assay, at a comparable sequencing depth. The improved resolution of the Visium HD array yields a larger number of clusters, identified via unsupervised clustering, that are well aligned with tissue morphology. To assess the spatial robustness of the assay, we quantified the abundance of canonical markers of normal colon epithelial cells in their expected location and adjacent cells, demonstrating that Visium HD has high transcript spatial localization accuracy. Together, these results demonstrate the high sensitivity, resolution, and accuracy of the Visium HD technology.

These features enabled us to perform an in depth analysis of a subset of three FFPE colon adenocarcinoma samples. The unsupervised clustering analysis of these samples allowed us to detect a broad range of cell types within each sample, however, the intra-patient heterogeneity made comparison between samples challenging. To ensure consistency in our cell type annotation and to increase our confidence in cell type calling, we generated a single cell reference dataset containing representative normal and diseased states. We then used this single cell dataset for deconvolution of Visium HD data from all of our samples. We adapted existing deconvolution methods that were designed based on Visium v2 to be performant with the dramatically increased number of ‘spots’ (now bins) that are present in Visium HD. Our analysis shows that per section unsupervised clustering analysis of Visium HD data yields similar cellular annotations for the main cell types found within tissue sections compared to the deconvolution method. While the inclusion of the single cell reference data provides an added benefit for a consistent cellular annotation strategy across multiple samples and identification of rare cell types, it is not a requirement for sample analysis using Visium HD.

The interactions of immune cells in the TME of CRC are poorly understood, hindering the development of new therapies^26^. For example, tumor-associated macrophages (TAMs) have been shown by several studies to exert pro-tumor activity; however, their role in CRC progression and the ability to predict disease outcomes based on macrophage infiltration are controversial. These controversial associations could be due to simultaneous accumulation of M1-like (pro-inflammatory) and M2-like (anti-inflammatory) macrophages and their spatial distribution within the tumor regions, leading to distinct functional activities within the TME^35,36^. It has been hypothesized that undifferentiated tumor cells at the invasion front, where the tumor tissue meets and interacts with the surrounding normal tissue, could polarize macrophages toward the M2-like phenotype (*SPP1*^+^ macrophages)^26^. Therefore, we wanted to explore the immune cell composition of the area immediately surrounding the tumor, to see what role immune cells were playing. In each sample, we identified distinct tumor cell types that were mapped to the morphological tumor regions in each tissue section, and applied a periphery analysis to interrogate areas within 50 µm of the tumor boundary. We observed a consistent enrichment of CAFs and macrophages in CRC tumor regions across all samples. CAFs are the most abundant non-immune cell types in the vicinity of CRC tumors, and TAMs are known to be the most prevalent immune cell types in the TME, recruited by cytokines released by tumor cells and CAFs. The presence of CAFs and *SPP1*+ macrophages are known to be highly correlated, and their presence is negatively correlated with lymphocyte infiltration and predict a poor patient survival^37^.

We next sought to understand the functional profiles of the macrophages identified in the tumor regions of our samples. Independent unsupervised clustering analysis of the gene signatures associated with the 8 µm bins labeled as macrophages found in tumor regions revealed two heterogeneous M2-like subpopulations, labeled as *SPP1*^+^ and *SELENOP*^+^ macrophages. These populations were specifically enriched within distinct spatial locations of the tumor, across two of the CRC tissue sections evaluated (only *SELENOP*^+^ macrophages were mapped within tumor regions in sample P5 CRC). *SELENOP*^+^ macrophages co-localized with tumor cell populations marked by expression of *REG* family genes, which are known to be highly expressed in CRC and associated with metastasis, advanced tumor stage and poor prognosis^38^. *SPP1^+^* macrophages co-localized with tumor cell populations marked by expression of *TGFBI,* which has been reported to be associated with poorer prognosis^39^. Previous scRNA-seq studies have shown that *SPP1^+^* macrophages are enriched in tumor tissue, exerting pro-tumor and pro-metastatic roles^9,26^. They may also regulate CAF through *TGFB1*, thereby promoting the secretion of MMPs and collagen to remodel the ECM, contributing to the resistance to PD-L1 blocking immunotherapy^37^. Pathway enrichment analysis of the upregulated gene expression profiles from both macrophage subpopulations revealed pathways consistent with pro-tumor activity. However, different pathways were dominant in each subpopulation, indicating that both subpopulations of macrophages identified here may exert pro-tumor effects, but do so by suppressing the immune response and contributing to tumor progression via different mechanisms. The resolution gains afforded by the increased density of the Visium HD arrays provided us with the ability to pinpoint the location of these subpopulations and the tumor cells they are interacting with, providing key insights into their behavior and the cellular dynamics of the TME. This is biologically meaningful because anti-PD-1 therapy is currently only effective for a minority of CRC patients, and disrupting the interactions of *SPP1*+ macrophages and CAFs has been proposed as a potential therapeutic strategy^26,37^.

The interesting dynamics we observed in the macrophage subpopulations led us to explore the immune cell populations in the tumors further, and we turned our attention to the adaptive immune response, specifically T cells. T cell infiltration into CRC tumors has long been associated with favorable outcomes, suggesting a possible role for immunoediting in controlling tumor growth^40,41^. Our analysis of Visium HD data at 8 µm bin size allowed us to clearly identify T cells in the microenvironment of the tumors we examined. However, we saw fewer T cells than anticipated based on the single cell data. Reanalyzing the data in this region using higher resolution (2 µm bins), enabled us to improve our capacity to identify T cells within the TME, since many were lost due to their small size and colocalization with other cells at 8 µm resolution. The 2 µm analysis also allowed us to improve accuracy, and pinpoint the location of the T cells in the tumor and surrounding areas, but even with this granular view we see that there is only a sparse presence of T cells in the TME. This resolution level flexibility was useful for answering specific biological questions, in this case the smaller bins provide critical insights for the analysis of T cells, but it was not necessary for the analysis of the larger macrophages. We anticipate that in the future more sophisticated analysis methods would be able to take full advantage of the 2 µm resolution data. Development of spatially aware methods that are able to include information from neighboring bins during unsupervised clustering would provide more accurate cell type annotations.

Finally, we wanted to explore the nature of the T cells we identified and better understand their function in the TME. Single cell TCR profiling allowed us to identify an expanded TCR clonotype within the more advanced cancer P5 CRC sample (stage IV-A, Table 1), suggesting an active immune response in this tumor. Using Xenium we mapped, for the first time, the location of clonally expanded T cells within the CRC TME, at single cell resolution. Xenium analysis revealed a co-localization of these clonally expanded T cells with cells expressing *CXCL9*, *CXCL10*, and *CXCL11* chemokines, which are known to attract cytotoxic T lymphocytes^34^. Consistent with this finding, these expanded T cells expressed cytotoxic genes, including *PRF1*, *GZMA*, and *GZMK*. Notably, macrophages within these regions emerged as the primary source of the *CXCL9*, *CXCL10*, and *CXCL11* expression, suggesting their contribution to T cell recruitment and potential anti-tumor activity. This finding suggests that despite the overall immunosuppressive environment in the microenvironment of these tumors, there are niches where anti-tumor immune responses may be taking place. This is an important observation as the balance between the pro- and anti-tumor macrophages influences tumor progression and response to therapy. Further research into the plasticity of macrophages and their ability to switch between these states could provide potential targets for therapeutic strategies in cancer treatment^42,43^.

High definition spatial technologies are providing increasingly granular insights into cellular behavior in the TME. Given the poor survival rates of many cancers, identification of better predictive prognostic biomarkers that can be used in clinical strategies are needed. The ability to visualize and analyze tumors using cutting edge spatial technologies not only enhances our comprehension of cancer biology but also guides the development of targeted therapies and has the potential to identify biomarkers that can meet this need. Our results highlight some of the insights that can be gleaned by studying immune cell populations in the TME with high definition, whole transcriptome, spatial technologies and pave the way for future studies that will fill gaps in our understanding of tumor evolution, progression, and provide insights for therapeutic advances.

## Supporting information

Supplemental Tables - Oliveira, Romero, Chung et al. 2024. HD spatial profiling of CRC.

Supplemental Material - Oliveira, Romero, Chung et al. 2024. HD spatial profiling of CRC.

## Data Availability

All datasets are available for download here: https://www.10xgenomics.com/products/visium-hd-spatial-gene-expression/dataset-human-crc

## Code Availability

Custom scripts used for this paper that are available on GitHub at: https://github.com/10XGenomics/HumanColonCancer_VisiumHD

## Acknowledgments

We would like to thank the Visium HD Development Team who built the Visium HD Spatial Gene Expression product used in this manuscript and the 10x Leadership Team for their support in this work. We also thank the members of the 10x Genomics microscopy, flow, sequencing, and histo-pathology core facilities.

## Author Contributions

M.O., J.R., and S.T. conceived the study; M.O., M.C., A.G., S.P., S.M., D.S., and D.P. performed experiments; M.O, J.R, M.C, S.W., and S.T. analyzed and interpreted data; M.O, J.P., M.C., A. D. G., and S.T. wrote the manuscript.

## Competing Interests

All authors are current or former employees or shareholders of 10x Genomics.

## Additional Information

Supplementary information can be found in an additional file.

## Methods

### Biomaterials

An overview of the experimental design is presented in Figure 1. We included samples from five patients with colon adenocarcinoma (two males, three females, ages 58-72, pre-treatment) in this study. From each patient, we included CRC FFPE blocks and NAT blocks for a subset of three patients (bringing the total to eight FFPE blocks), and we obtained paired fresh frozen dissociated tumor cells (DTCs), essential for immune profiling analysis, alongside the three selected FFPE blocks used for spatial profiling (Table 1).

### Tissue sectioning

5 µm sections were taken from the FFPE tissue blocks with a microtome (Epredia HM355S).

Sections were adjacent or near-adjacent (within 5-10 µm of each other). Sectioning followed the Xenium In Situ for FFPE - Tissue Preparation Guide (CG000578, Rev C) for the Xenium workflow, or the Visium CytAssist Spatial Gene Expression for FFPE – Tissue Preparation Guide (CG000518, Rev C) for the Visium workflows.

### Visium HD Spatial Gene Expression

Like first-generation Visium assays, the Visium HD assay is compatible with FFPE-embedded tissues and H&E / IF staining. However, the Visium HD workflow requires new HD slides that require thawing, washes, and equilibration in appropriate buffers. Visium HD slides also feature high-resolution fiducials for subpixel image alignment, and a dispensing pad and spacer for CytAssist compatibility. We first placed FFPE tissue sections on plain glass slides for deparaffinization, H&E staining and imaging following the Visium HD FFPE Tissue Preparation Handbook (CG000684). Probe hybridization, probe ligation, slide preparation, probe release, extension, library construction, and sequencing followed the Visium HD Spatial Gene Expression Reagent Kits User Guide (CG000685). Sequencing was performed on an Illumina NovaSeq 6000 with paired-end reads (43 cycles Read 1, 10 cycles i7, 10 cycles i5, 50 cycles Read 2). We used Space Ranger v3.0 to map FASTQ files to the human reference, detect the tissue section, align the sequencing data to the microscope image and the CytAssist image, and output gene-barcode matrices for further analysis.

### Visium CytAssist Spatial Gene Expression for FFPE

We ran Visium CytAssist Spatial Gene Expression for FFPE (“Visium v2”) on a subset of samples to demonstrate technological improvements of Visium HD. 5 µm FFPE serial sections were placed on standard glass slides and H&E-stained following the Demonstrated Protocol Visium CytAssist Spatial Gene Expression for FFPE – Deparaffinization, H&E Staining, Imaging & Decrosslinking (CG000520). Imaging was performed on the slide scanner Olympus V200 at 20x magnification. The CytAssist instrument was used to facilitate the transfer of transcriptomic probes from the standard glass slide to the Visium CytAssist Spatial Gene Expression Slide, v2, 11 mm capture area. Probe hybridization, probe ligation, release, extension, pre-amplification, and library preparation followed the Visium CytAssist Spatial Gene Expression Reagent Kits User Guide (CG000495). Sequencing was performed on the NovaSeq 6000 (28 cycles Read 1, 10 cycles i7, 10 cycles i5, 90 cycles Read 2). Flow cells were demultiplexed using the mkfastq command in Space Ranger (v3.0). FASTQs were aligned to the human (GRCh38) reference with Space Ranger v3.0.

### Chromium Single Cell Gene Expression Flex

We collected Chromium Single Cell Gene Expression Flex data (10x Genomics) to create a sample-specific annotated reference atlas. Cells were dissociated from 50 µm FFPE curls from CRC tissue samples (n=5 patients) and a subset of paired normal adjacent tissues (n = 3 patients) using the Demonstrated Protocol for Isolation of Cells from FFPE Tissue Sections for Chromium Fixed RNA Profiling (CG000632). The FFPE curls used for this analysis were serial to the sections used for the Visium HD assay, to ensure that the single cell reference atlas would closely represent the tissue analyzed with the spatial analysis. Flex library preparation followed the Chromium Fixed RNA Profiling for Multiplexed Samples User Guide (CG000527, RevD). Each sample was hybridized with a unique probe barcode in an individual reaction. After hybridization, samples were multiplexed in equal cell proportions, washed, and processed collectively, across multiple GEM lanes as technical replicates, using the Chromium X instrument. Libraries were sequenced on an Illumina NovaSeq 6000 with paired-end dual-indexing (28 cycles Read 1, 10 cycles i7, 10 cycles i5, 90 cycles Read 2). Flow cells were demultiplexed using the mkfastq command in Cell Ranger (v8.0.0). Cell Ranger v8.0.0 (10x Genomics) was used to align reads in FASTQ format to the human probe set and reference genome (GRCh38), producing feature-barcode matrices for analysis. Each GEM well was processed using a separate instance of the cellranger multi pipeline. Finally, we aggregated matrices across all eight tissue blocks with the cellranger aggr pipeline.

After sequencing, we aggregated the single cell data and analyzed it as a combined dataset from all patients. To build the atlas, we used Seurat v5^44^ to import the H5 file produced by the cellranger aggr pipeline. We filtered the data on quality control metrics including the number of features, the number of UMIs per barcode, and the percentage of mitochondrial UMIs. We retained barcodes with fewer than 25% mitochondrial UMIs, as tumors are expected to have higher mitochondrial expression.

We then plotted the distribution of UMIs and genes per barcode, excluding the top and bottom 2.5% of the distribution to account for outliers. Given the dataset’s large size, we adopted the sketch-based analysis approach in Seurat (https://satijalab.org/seurat/articles/seurat5_sketch_analysis), sampling 15% of the entire dataset (∼37,000 cells) for downstream analysis. Using Seurat, we identified variable features, scaled the data, performed PCA, identified neighbors, and conducted graph-based clustering (retaining 25 principal components (PCs) with a resolution parameter of 0.6). We then applied differential gene expression (DGE) analysis to identify marker genes within these clusters. We manually classified the graph-based clusters into ten broad cell types, denoted as level 1 annotations. For level 2 annotations, we repeated the clustering process within each level 1 cluster, maintaining 25 PCs but adjusting the resolution parameter to 0.1 to prevent over-splitting. We identified marker genes through DGE and annotated cell types manually. After completing the annotations on the subsampled data, we extended the annotations to the entire single cell dataset.

When single cell datasets were segregated per patient, different proportions of cell types as well as distinct types of tumor cells were revealed that were differently enriched per patient, highlighting the inter-patient heterogeneity of CRC samples. We further validated these findings by performing a differential expression analysis, which revealed patient specific tumor signatures (Supplemental Figures 3 and 4).

### Single Cell Immune Profiling

Single cell immune profiling relies on poly-A-based 5’ mRNA capture and targeted sequencing and assembly of the V(D)J regions, therefore fresh frozen dissociated tumor cells (DTCs) are necessary. We used DTCs that were obtained from the same tumor samples as our FFPE blocks to obtain TCR profiling data on three samples (Table 1). Cryovials were thawed in a water bath for 2–3 minutes at 37°C with gentle shaking. Thawed cells were transferred into a 15 mL centrifuge tube containing 9 mL of pre-warmed complete growth medium (RPMI + 10% FBS) and the centrifuged at 150 rcf for 10 minutes at room temperature using a swinging bucket rotor. All but ∼200 μL of the supernatant was removed and then the pellet was resuspended in this remaining 200 μL, followed by the addition of 200 μL of complete media. Cells were strained using a 40 μm Flowmi cell strained and transferred to a 1.5 mL Eppendorf tube before being counted on a Cellaca MX using AOPI. 5 μL of CD3 BV510 stain and 2.5uL of CD45 APC was added to the cell suspension and it was incubated on ice for 30 minutes in the dark. Cells were then centrifuged at 300 rcf for 10 minutes at 4°C. The supernatant was removed without disturbing the cell pellet and 1 mL of chilled PBS + 10% FBS added to the tube with gentle mixing. Cells were centrifuged at 300 rcf for 10 minutes at 4°C and the pellet resuspended in 250 μL of chilled PBS + 10% FBS. Cells were then sorted on a SONY MA900 cell sorter. Live single DTCs were gated using forward scatter (relative size) and back scatter (relative granularity) and 7AAD for live/dead cell discrimination. CD45 and CD3 were used to identify T cells. Sorts were performed on a 100 µm nozzle at 20 psi sheath pressure. The sample pressure was set to low-medium with a consistent event rate maintained throughout the sort. Cells were sorted into 30 μL 10% FBS in PBS, in a pre-coated 1.5 mL Eppendorf tube and then directly loaded onto the 10x chip for generation of 5′ gene expression and TCR libraries following the Chromium Next GEM Single Cell 5′ Reagent Kits v2 (Dual Index) User Guide (CG000331) on the Chromium X instrument. Libraries were sequenced on an Illumina NovaSeq 6000 with paired-end dual-indexing (28 cycles Read 1, 10 cycles i7, 10 cycles i5, 90 cycles Read 2). Flow cells were demultiplexed using the mkfastq command in Cell Ranger (v8.0.0). We used Cell Ranger v8.0.0 (10x Genomics) to process FASTQ data for both gene expression and TCR libraries. The cellranger count pipeline was used to align 5’ reads to the reference genome (GRCh38) and output gene-barcode matrices that could be used to cluster cells and run differential gene expression. We used the cellranger vdj pipeline to assemble TCR sequences and group cells into clonotypes based on their CDR3 sequences.

### Xenium In Situ

We ran Xenium In Situ on sample P1 CRC, P2 CRC, and P5 CRC to validate Visium HD results, providing a direct comparison for gene sensitivity, and to map a clonal expansion of T-cells detected in sample P5 CRC. The 10x Genomics Xenium Human Colon Gene Expression Panel (322 genes) was supplemented with an additional 100 genes chosen to characterize the TME (for the complete gene list see Supplemental Table 1). Three genes, each with unique probesets, were designed for the two alpha and one beta chains of the TCR clonotype identified from the Single Cell Immune Profiling data in sample P5 CRC. The other 97 genes included markers that were enriched in different subsets of tumor cells from the single cell reference atlas and from the Visium HD data. They include marker genes for macrophages, neutrophils, cancer-associated fibroblasts, granulocytes, chemokines, and other TME markers. The panel was designed using Xenium Panel Designer following the guidance in the Xenium Add-on Panel Design Technical Note (CG000643, RevB).

During the Xenium Analyzer workflow, the Xenium In Situ Cell Segmentation Solution uses a Multi-Tissue Stain Mix to identify cell boundaries (cell segmentation) in an automated fashion. The solution includes antibodies for labeling membranes, antibodies for labeling cell interiors, a universal interior label for ribosomal RNA, and a DAPI stain for nuclei. Probe hybridization, ligation, amplification, cell segmentation staining, and autofluorescence quenching followed the Xenium In Situ Gene Expression with Morphology-based Cell Segmentation Staining User Guide (CG000749). The Xenium Onboard Analysis pipeline v2.0.0 (10x Genomics) was run directly on the instrument for imaging processing, cell segmentation, image registration, decoding, deduplication, and secondary analysis.

### Design of TCR clonotype probes

Custom probes for Xenium were developed to target three CDR3 sequences identified by VDJ sequencing. One 40 bp probe was designed for each CDR3, centered on the CDR3 with some overhang into the adjacent framework regions. These probes were designed following the specifications in the [Species Standalone Custom and Advanced Custom Panel Design for Xenium In Situ Technical Note (CG000683, Rev C)](https://www.10xgenomics.com/support/in-situ-gene-expression/documentation/steps/panel-design/species-standalone-custom-and-advanced-custom-panel-design-for-xenium-in-situ). Probe sequences are shown in Table 2.

**Table 2.**
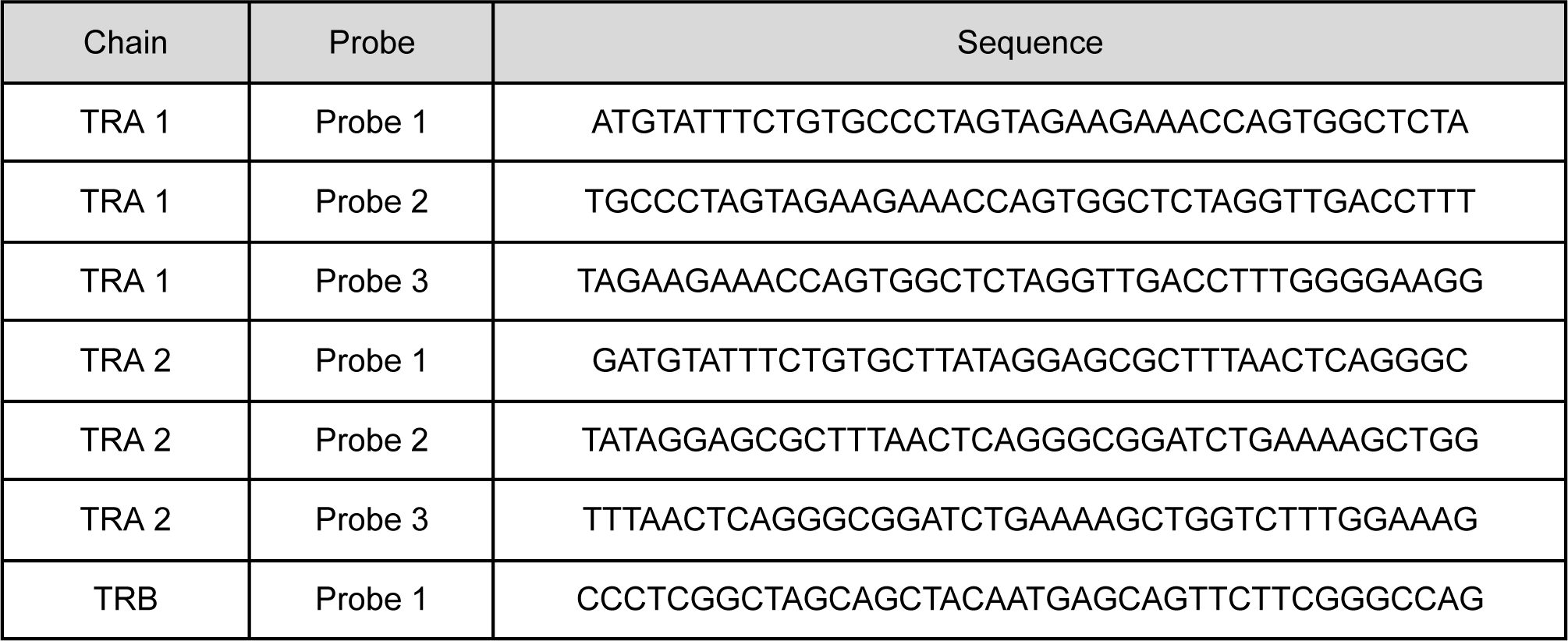
TCR Clonotype Xenium Probes.

### Visium v2 to HD sensitivity comparison

We assessed the sensitivity of Visium HD compared to Visium v2 on a gene-by-gene basis. Matched areas were manually selected in Loupe and probe-barcode matrices from each dataset, generated by Space Ranger, were imported into Seurat v5^44^ using the read10xh5() function. We utilized the ggplot2 R package to graph per-probe UMI counts. The data were displayed on a Log10 +1 scale, with Spearman correlations calculated as r^2^. Our comparison spanned all probes across the entire transcriptome and specifically focused on probes crossing an exon-exon splice junction. This latter comparison helps exclude most probes that could target genomic DNA (gDNA) or be susceptible to off-target effects. For more details on genomic DNA see [Visium CytAssist Spatial Gene Expression for FFPE: Robust Data Analysis with Minimal Impact of Genomic DNA](https://cdn.10xgenomics.com/image/upload/v1666737555/support-documents/CG000605_GenomicDNAinVisiumCytAssistSpatialAssay_RevA.pdf) Technical Note (CG000605, Rev A).

### Spatial accuracy

To measure spatial accuracy of mRNA detection, we identified morphologically distinct regions of interest (ROIs) and then pinpointed marker genes unique to each ROI. These marker genes should be expressed only in squares directly beneath their corresponding ROI. Using QuPath v0.4.4^45^, we selected four ROIs within normal colon mucosal glands (hereinafter referred to as “source masks”) and areas of adjacent muscularis mucosae (“adjacent masks”), choosing three goblet cell marker genes (*CLCA1*, *FCGBP*, *MUC2*). We mapped the locations of all transcripts for these marker genes and calculated the proportion of accurately localized transcripts for each of the four ROIs. For the remaining transcripts, we determined the Euclidean distance from the edge of the nearest source mask to the square of transcript detection. Additionally, we calculated the densities of the marker genes within both the source and adjacent masks for each ROI.

### Spot deconvolution

Spot deconvolution was used to classify and label bins with cell types derived from the single cell atlas. We ran spacexr^46^ using doublet mode, which restricts any given bin to at most two cell types. Spacexr is an open source package, and we modified the code to improve runtime due to the increased ‘spot’ number possible with Visium HD. In Visium HD, there is a maximum upper limit of 11,222,500 barcoded 2 μm bins within a 6.5 x 6.5 mm capture area, whereas Visium v2 has a maximum of approximately 14,000 barcoded 55 μm spots within an 11 x 11 mm capture area (or 5,000 barcoded 55 μm spots within an 6.5 x 6.5 mm capture area). The minimum UMI threshold was 100.

### Characterizing the tumor periphery

Using the deconvolution results, we used a custom script to identify the peripheral region (up to 50 µms from the tumor). Briefly, for each tumor bin, we selected all other bins within 50 µm that were not classified as tumor. We also removed any tumor bin that had less than 25 tumor neighbors, to reduce isolated tumor bins in the tissue. Then, we obtained the cell type proportions in the peripheral region, and compared those proportions to cell types observed in the rest of the tissue.

### Immune density analysis

To map the locations of specific immune cell types in the CRC samples, we used the coordinates for each bin and their labels provided by deconvolution. We counted only bins that were labeled as singlets. We used a 2D kernel density estimation to select regions enriched in a given cell type. Density values were scaled to a maximum of 1.

### Distance / local / regional analysis

After identifying regions enriched with immune cells, we selected the top three regions exhibiting the highest macrophage density (within bins categorized as tumor tissue) and delineated these areas as Regions of Interest (ROIs) with a radius of 350 µm. Additionally, we identified a “cold” region lacking immune infiltration. To identify genes associated with elevated immune cell density, we used differential gene expression (DGE) analysis. Subsequent enrichment analysis of differentially expressed genes (ranked by log_2_ fold change) used the Hallmark gene sets linked to specific biological pathways^47^, https://www.gsea-msigdb.org/gsea/msigdb/human/collections.jsp).

### Nuclei Segmentation

To segment nuclei from the H&E images and assign 2 µm bins to the identified nuclei, we followed the analysis guide “Nuclei Segmentation and Custom Binning of Visium HD Gene Expression Data’’ (https://www.10xgenomics.com/analysis-guides/segmentation-visium-hd). The segmentation procedure was run on the full section using stardist^48^. We used affine transformations to preserve the segmentation polygons when subsetting the image to specific regions of interest. Once the 2 µm bins were assigned to the corresponding nuclei polygons, the data was transformed to create a gene by nuclei UMI count matrix for further processing.

## Notes

https://www.10xgenomics.com/products/visium-hd-spatial-gene-expression/dataset-human-crc

https://github.com/10XGenomics/HumanColonCancer_VisiumHD

## References

1. Xi, Y. & Xu, P. Global colorectal cancer burden in 2020 and projections to 2040. Transl. Oncol. 14, 101174 (2021).

2. Hossain, M. S. et al. Colorectal Cancer: A Review of Carcinogenesis, Global Epidemiology, Current Challenges, Risk Factors, Preventive and Treatment Strategies. Cancers 14, 1732 (2022).

3. O’Connell, J. B., Maggard, M. A. & Ko, C. Y. Colon Cancer Survival Rates With the New American Joint Committee on Cancer Sixth Edition Staging. JNCI J. Natl. Cancer Inst. 96, 1420–1425 (2004).

4. Wang, W. et al. Molecular subtyping of colorectal cancer: Recent progress, new challenges and emerging opportunities. Semin. Cancer Biol. 55, 37–52 (2019).

5. Guinney, J. et al. The consensus molecular subtypes of colorectal cancer. Nat. Med. 21, 1350–1356 (2015).

6. Singh, M. P., Rai, S., Pandey, A., Singh, N. K. & Srivastava, S. Molecular subtypes of colorectal cancer: An emerging therapeutic opportunity for personalized medicine. Genes Dis. 8, 133–145 (2021).

7. Sawayama, H., Miyamoto, Y., Ogawa, K., Yoshida, N. & Baba, H. Investigation of colorectal cancer in accordance with consensus molecular subtype classification. Ann. Gastroenterol. Surg. 4, 528–539 (2020).

8. Wen, R. et al. Single-cell sequencing technology in colorectal cancer: a new technology to disclose the tumor heterogeneity and target precise treatment. Front. Immunol. 14, (2023).

9. Zhang, L. et al. Single-Cell Analyses Inform Mechanisms of Myeloid-Targeted Therapies in Colon Cancer. Cell 181, 442–459.e29 (2020).

10. Pelka, K. et al. Spatially organized multicellular immune hubs in human colorectal cancer. Cell 184, 4734–4752.e20 (2021).

11. Becker, W. R. et al. Single-cell analyses define a continuum of cell state and composition changes in the malignant transformation of polyps to colorectal cancer. Nat. Genet. 54, 985–995 (2022).

12. Avraham-Davidi, I. et al. Integrative single cell and spatial transcriptomics of colorectal cancer reveals multicellular functional units that support tumor progression. 2022.10.02.508492 Preprint at 10.1101/2022.10.02.508492 (2022).

13. Joanito, I. et al. Single-cell and bulk transcriptome sequencing identifies two epithelial tumor cell states and refines the consensus molecular classification of colorectal cancer. Nat. Genet. 54, 963–975 (2022).

14. Khaliq, A. M. et al. Refining colorectal cancer classification and clinical stratification through a single-cell atlas. Genome Biol. 23, 113 (2022).

15. Li, H. et al. Reference component analysis of single-cell transcriptomes elucidates cellular heterogeneity in human colorectal tumors. Nat. Genet. 49, 708–718 (2017).

16. Lee, H.-O. et al. Lineage-dependent gene expression programs influence the immune landscape of colorectal cancer. Nat. Genet. 52, 594–603 (2020).

17. Cho, C.-S. et al. Microscopic examination of spatial transcriptome using Seq-Scope. Cell 184, 3559–3572.e22 (2021).

18. Poovathingal, S. et al. Nova-ST: Nano-Patterned Ultra-Dense platform for spatial transcriptomics. 2024.02.22.581576 Preprint at 10.1101/2024.02.22.581576 (2024).

19. Schott, M. et al. Open-ST: High-resolution spatial transcriptomics in 3D. 2023.12.22.572554 Preprint at 10.1101/2023.12.22.572554 (2023).

20. Vickovic, S. et al. High-definition spatial transcriptomics for in situ tissue profiling. Nat. Methods 16, 987–990 (2019).

21. Liu, Y. et al. High-Spatial-Resolution Multi-Omics Sequencing via Deterministic Barcoding in Tissue. Cell 183, 1665–1681.e18 (2020).

22. Fu, X. et al. Polony gels enable amplifiable DNA stamping and spatial transcriptomics of chronic pain. Cell 185, 4621–4633.e17 (2022).

23. Lee, Y. et al. XYZeq: Spatially resolved single-cell RNA sequencing reveals expression heterogeneity in the tumor microenvironment. Sci. Adv. 7, eabg4755 (2021).

24. Peng, Z., Ye, M., Ding, H., Feng, Z. & Hu, K. Spatial transcriptomics atlas reveals the crosstalk between cancer-associated fibroblasts and tumor microenvironment components in colorectal cancer. J. Transl. Med. 20, 302 (2022).

25. Liu, H.-T. et al. Spatially resolved transcriptomics revealed local invasion-related genes in colorectal cancer. Front. Oncol. 13, (2023).

26. Ozato, Y. et al. Spatial and single-cell transcriptomics decipher the cellular environment containing HLA-G+ cancer cells and SPP1+ macrophages in colorectal cancer. Cell Rep. 42, (2023).

27. Wang, F. et al. Single-cell and spatial transcriptome analysis reveals the cellular heterogeneity of liver metastatic colorectal cancer. Sci. Adv. 9, eadf5464 (2023).

28. Tian, L., Chen, F. & Macosko, E. Z. The expanding vistas of spatial transcriptomics. Nat. Biotechnol. 41, 773–782 (2023).

29. Du, M. R. M. et al. Spotlight on 10x Visium: a multi-sample protocol comparison of spatial technologies. 2024.03.13.584910 Preprint at 10.1101/2024.03.13.584910 (2024).

30. Khalaf, K. et al. Aspects of the Tumor Microenvironment Involved in Immune Resistance and Drug Resistance. Front. Immunol. 12, (2021).

31. Ma, S.-X., Li, L., Cai, H., Guo, T.-K. & Zhang, L.-S. Therapeutic challenge for immunotherapy targeting cold colorectal cancer: A narrative review. World J. Clin. Oncol. 14, 81–88 (2023).

32. Janesick, A. et al. High resolution mapping of the tumor microenvironment using integrated single-cell, spatial and in situ analysis. Nat. Commun. 14, 8353 (2023).

33. Tian, X. et al. Expression of CD147 and matrix metalloproteinase-11 in colorectal cancer and their relationship to clinicopathological features. J. Transl. Med. 13, 337 (2015).

34. Tokunaga, R. et al. CXCL9, CXCL10, CXCL11/CXCR3 axis for immune activation – A target for novel cancer therapy. Cancer Treat. Rev. 63, 40–47 (2018).

35. Zhang, C., Yang, M. & Ericsson, A. C. Function of Macrophages in Disease: Current Understanding on Molecular Mechanisms. Front. Immunol. 12, (2021).

36. Jahandideh, A. et al. Macrophage’s role in solid tumors: two edges of a sword. Cancer Cell Int. 23, 150 (2023).

37. Qi, J. et al. Single-cell and spatial analysis reveal interaction of FAP+ fibroblasts and SPP1+ macrophages in colorectal cancer. Nat. Commun. 13, 1742 (2022).

38. Zhou, M. et al. N6-methyladenosine modification of REG1α facilitates colorectal cancer progression via β-catenin/MYC/LDHA axis mediated glycolytic reprogramming. Cell Death Dis. 14, 1–16 (2023).

39. Ma, C. et al. Extracellular matrix protein βig-h3/TGFBI promotes metastasis of colon cancer by enhancing cell extravasation. Genes Dev. 22, 308–321 (2008).

40. Yang, W. et al. T-cell infiltration and its regulatory mechanisms in cancers: insights at single-cell resolution. J. Exp. Clin. Cancer Res. 43, 38 (2024).

41. Teng, M. W. L., Galon, J., Fridman, W.-H. & Smyth, M. J. From mice to humans: developments in cancer immunoediting. J. Clin. Invest. 125, 3338–3346 (2015).

42. Ganesh, K. et al. Immunotherapy in colorectal cancer: rationale, challenges and potential. Nat. Rev. Gastroenterol. Hepatol. 16, 361–375 (2019).

43. Goossens, P. et al. Membrane Cholesterol Efflux Drives Tumor-Associated Macrophage Reprogramming and Tumor Progression. Cell Metab. 29, 1376–1389.e4 (2019).

44. Hao, Y. et al. Dictionary learning for integrative, multimodal and scalable single-cell analysis. Nat. Biotechnol. 42, 293–304 (2024).

45. Bankhead, P. et al. QuPath: Open source software for digital pathology image analysis. Sci. Rep. 7, 16878 (2017).

46. Cable, D. M. et al. Robust decomposition of cell type mixtures in spatial transcriptomics. Nat. Biotechnol. 40, 517–526 (2022).

47. Liberzon, A. et al. The Molecular Signatures Database Hallmark Gene Set Collection. Cell Syst. 1, 417–425 (2015).

48. Weigert, M. & Schmidt, U. Nuclei Instance Segmentation and Classification in Histopathology Images with Stardist. in 2022 IEEE International Symposium on Biomedical Imaging Challenges (ISBIC) 1–4 (IEEE, Kolkata, India, 2022). doi:10.1109/ISBIC56247.2022.9854534.

