## Supplemental Material - Oliveira, Romero, Chung et al. 2024. HD spatial profiling of CRC. for "Characterization of immune cell populations in the tumor microenvironment of colorectal cancer using high definition spatial profiling"

#### Supplemental Figures

##### Supplemental Figure 1

**A**

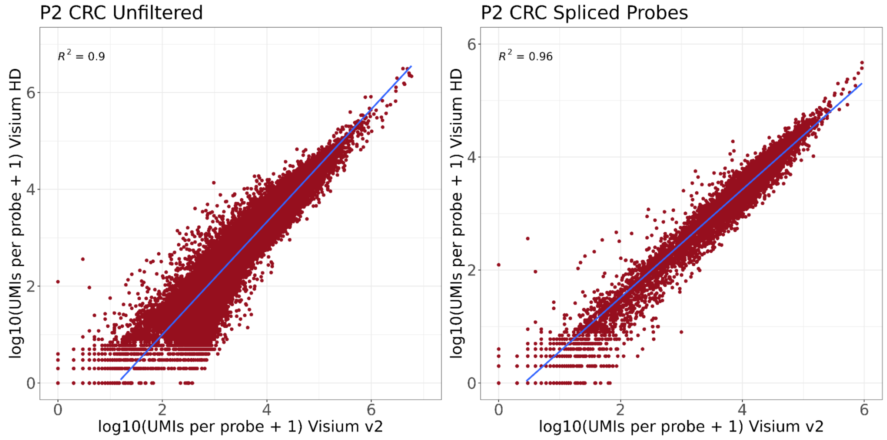


**B**

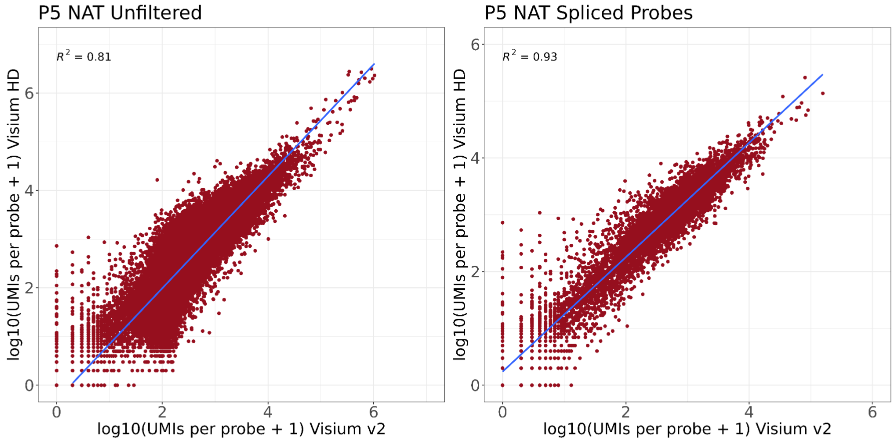


**Supplemental Figure 1. Sensitivity comparisons between Visium v2 and Visium HD performed on serial sections of normal and colon cancer samples.**
Comparisons show strong correlation between UMI counts from all probes (left, unfiltered), and the probes that only span spliced gene target regions (right, spliced probes), obtained from each assay, highlighting comparable sensitivity between assays.
A. CRC sample (P2 CRC). B. Normal colon mucosa sample (P5 NAT).

### Supplemental Figure 2

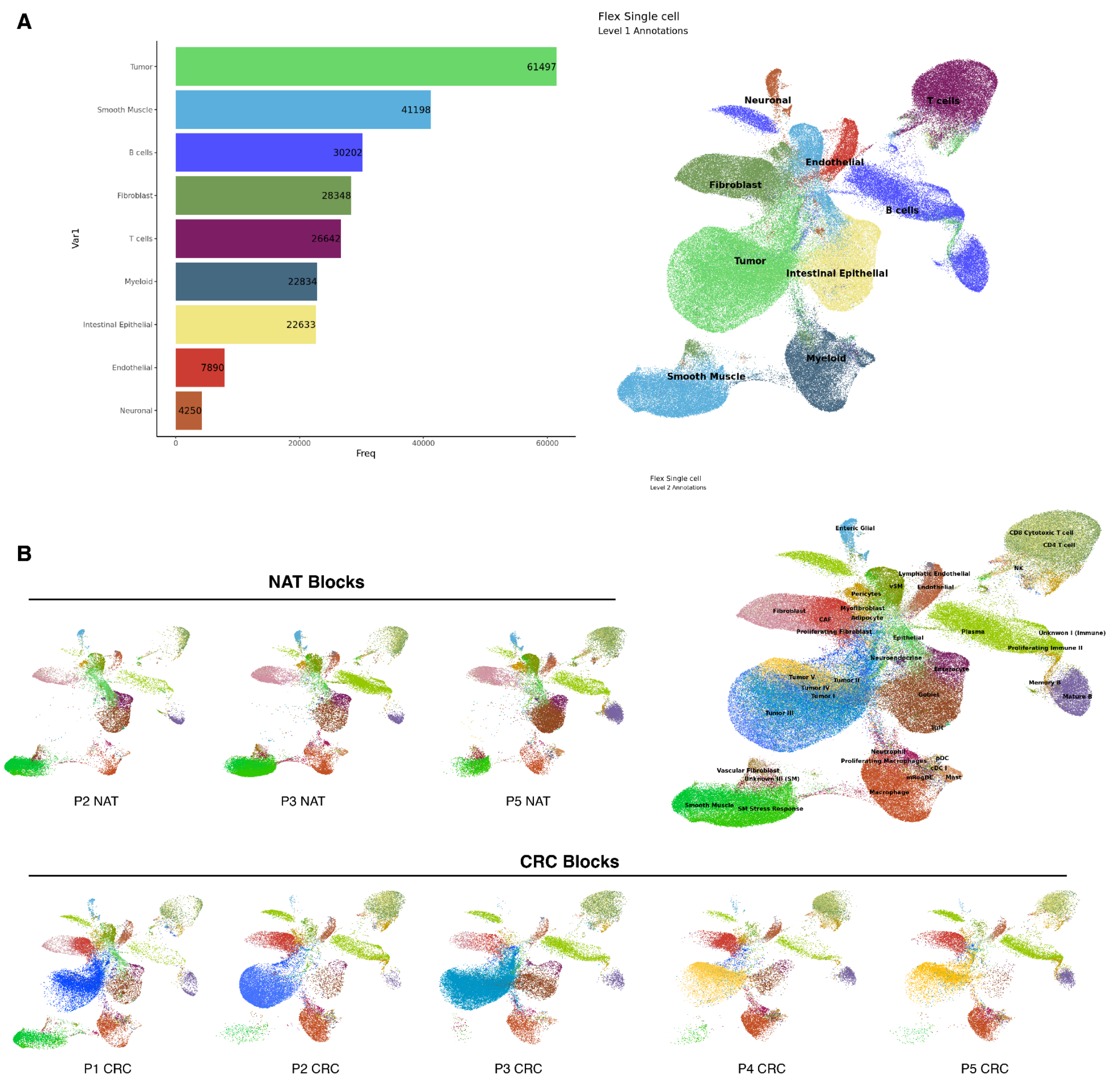


**Supplemental Figure 2: Cell type annotation of the sample-specific single cell reference atlas. A.** Left: Bar plot showing frequency of distinct cell types across the single cell data set composed of 5 CRC sections and 3 NAT sections. Annotations were based on published gene markers. Right: UMAP plot showing level 1 cell type annotations in the single cell dataset. Endothelial cells (7,890 cells), fibroblasts (28,348 cells), intestinal epithelial cells (22,633 cells), myeloid (22,834 cells), B cells (30,202 cells), neuronal cells (4,250 cells), smooth muscle cells (41,198 cells), T cells (26,642 cells), and tumor cells (61,497 cells).

**B.** Top left and bottom row: UMAP plots showing cell type annotations in individual samples after further sub-clustering analysis of the single cell data. Top right: UMAP plot showing level 2 cell type annotations in the single cell revealing 39 clusters: endothelial cells (3 clusters), fibroblasts (6 clusters), intestinal epithelial cells (3 clusters), myeloid (7 clusters), B cells (3 clusters), intestinal glial cells (1 cluster), smooth muscle cells (5 clusters), T cells (5 clusters), and tumor cells (5 clusters).

Supplemental Figure 3

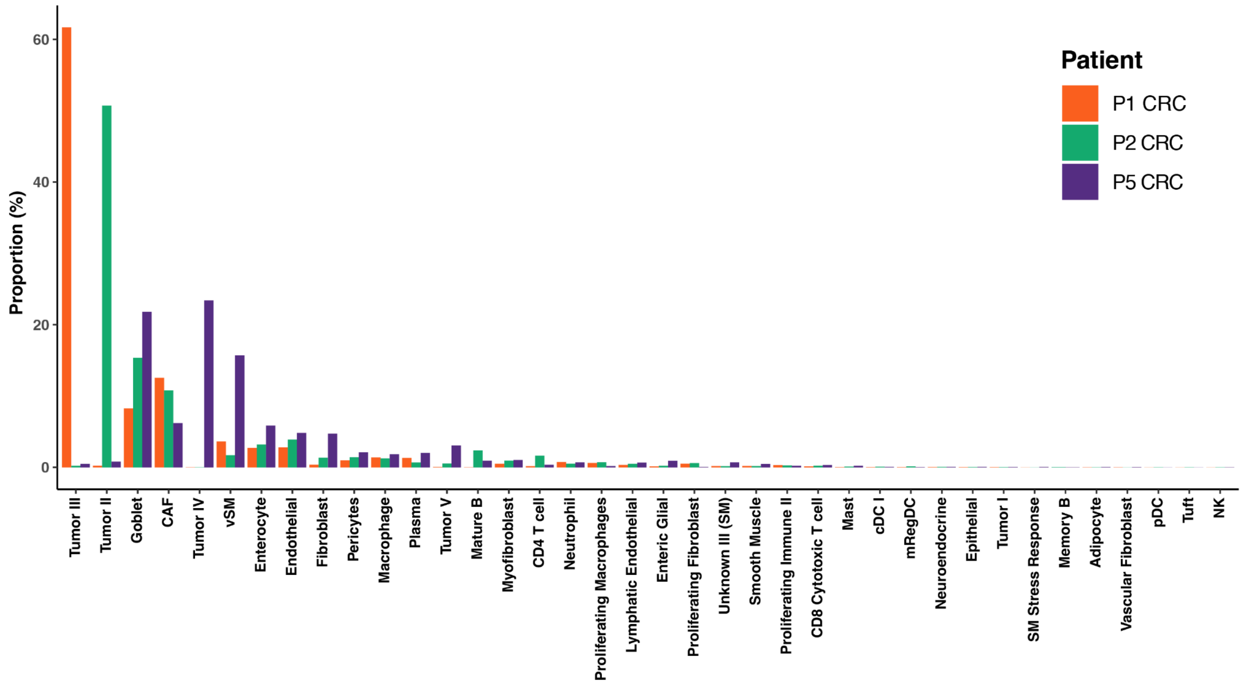


**Supplemental Figure 3:** Proportion of cell types identified after spot deconvolution of the Visium HD data, annotated using the level 2 cell type annotations from the combined single cell reference dataset shown in Supplemental Figure 2.

### Supplemental Figure 4

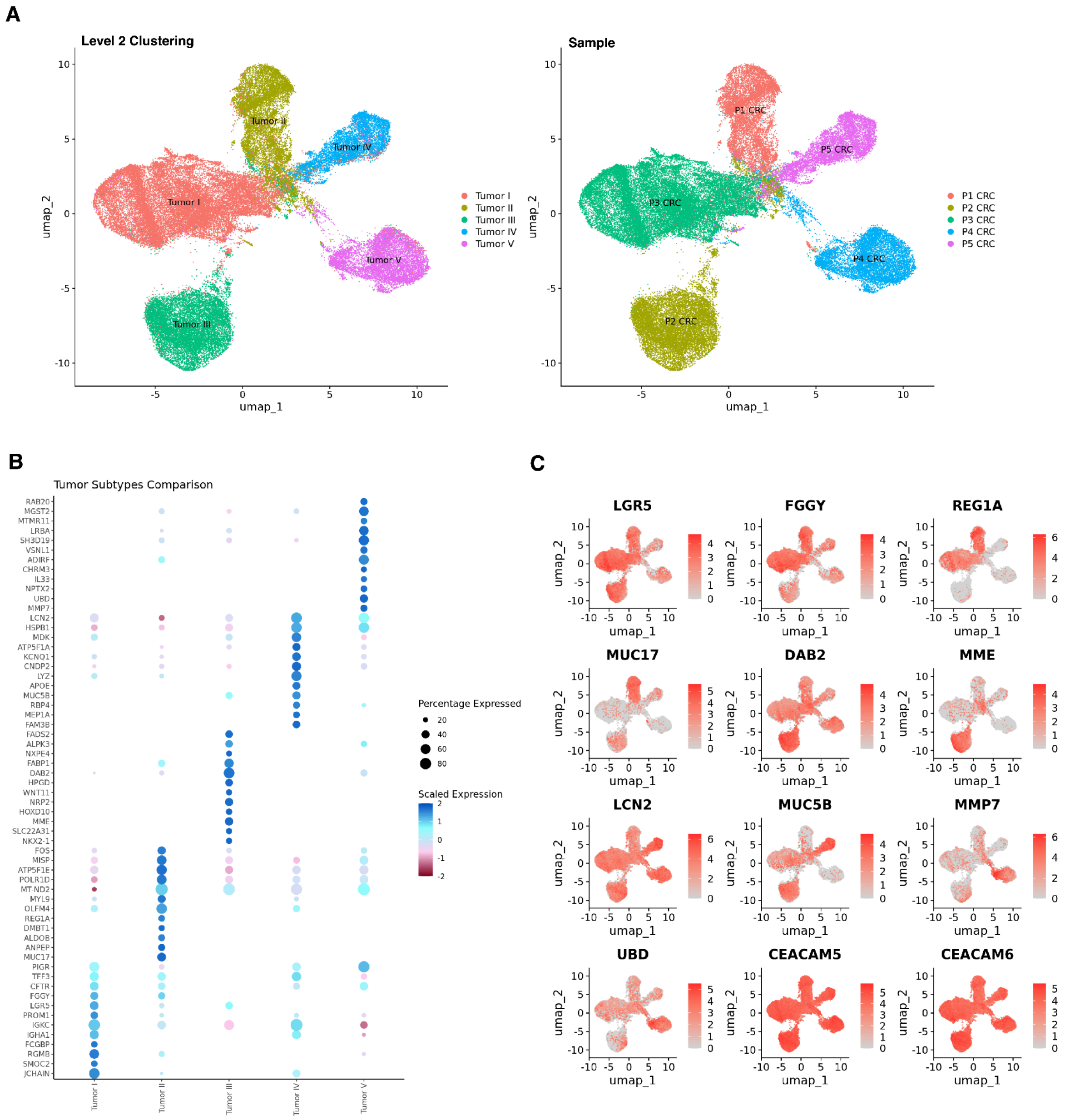


**Supplemental Figure 4: Single cell analysis of tumor subpopulations in CRC (n=5) samples reveal tumor heterogeneity. A.** UMAP plot with tumor cells colored by cluster (left) and colored by sample identifier (right). **B.** Dot plot displaying the scaled expression of the top differentially expressed genes across the 5 tumor subpopulations. **C.** UMAP plot colored by log normalized UMI counts of top differentially expressed genes.

Supplemental Figure 5

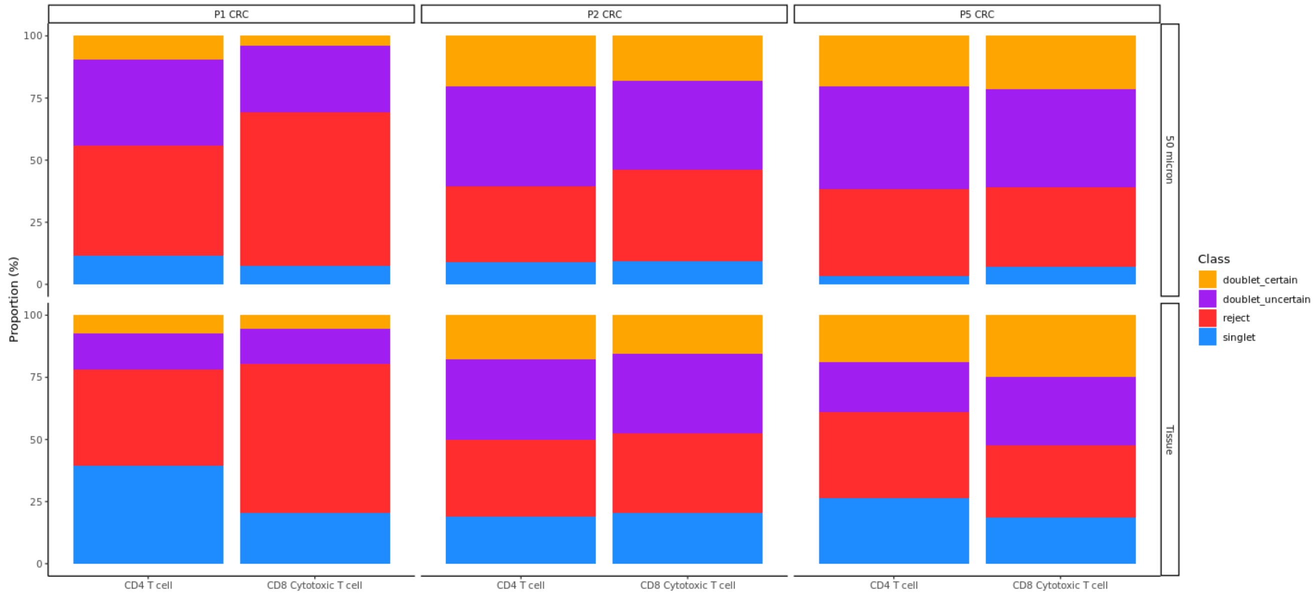


**Supplemental Figure 5. Deconvolution class for 8 μm bins labeled as T cells.** Barplots denoting the deconvolved class for CD4 T cells and CD8 cytotoxic T cells in the 50 micron TME and rest of the tissue for each patient.

Supplemental Figure 6

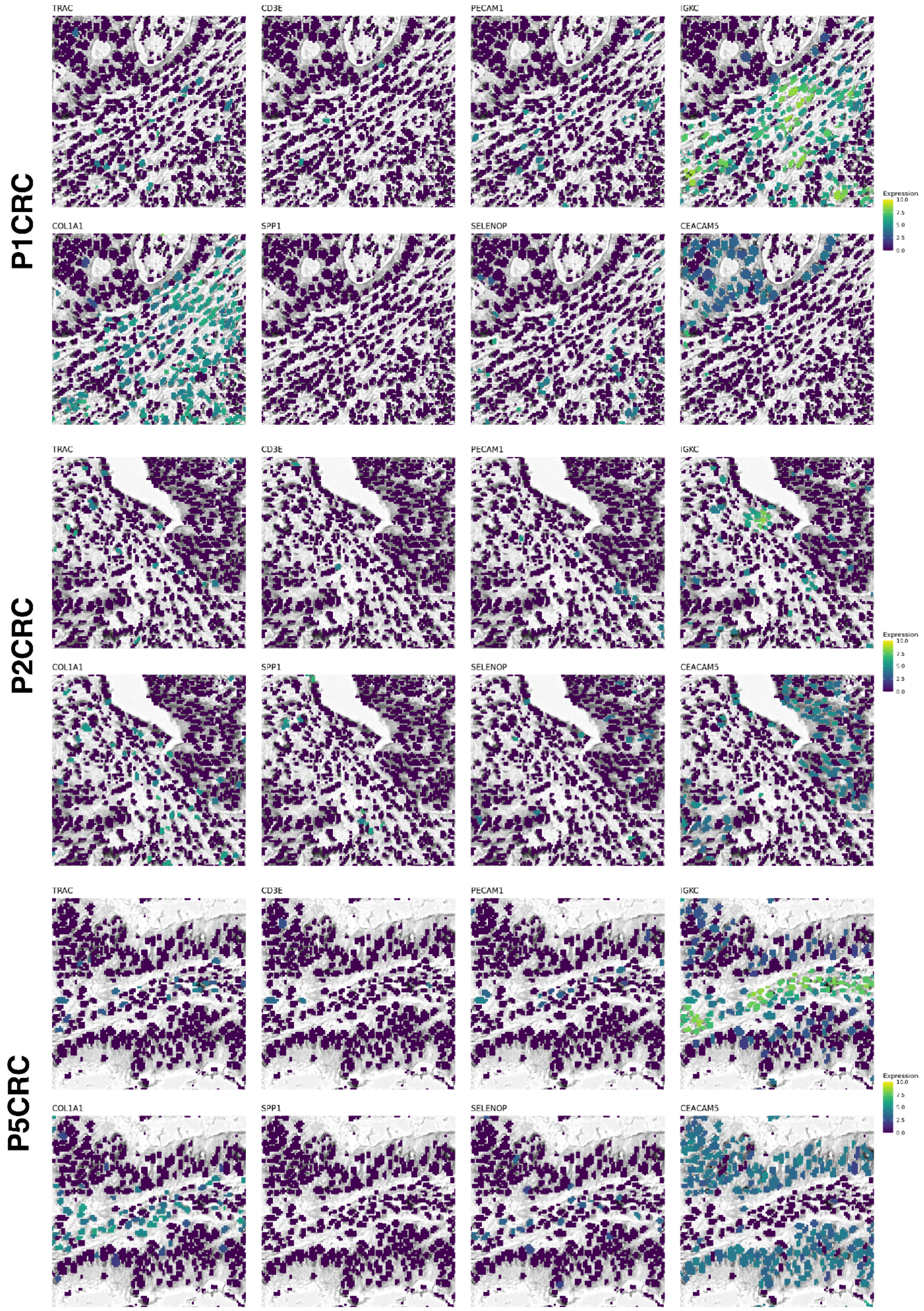


**Supplemental Figure 6:** Expression of *TRAC* (T cell), *CD3E* (T cell), *PECAM1* (Endothelial), *IGKC* (Plasma), *COL1A1* (CAF), *SPP1* (Macrophage), *SELENOP* (Macrophage) and *CEACAM5* (Tumor) in the segmented nuclei for each patient. UMI counts were grouped by 2 micron bins located within each segmented nuclei.

### Supplemental Figure 7

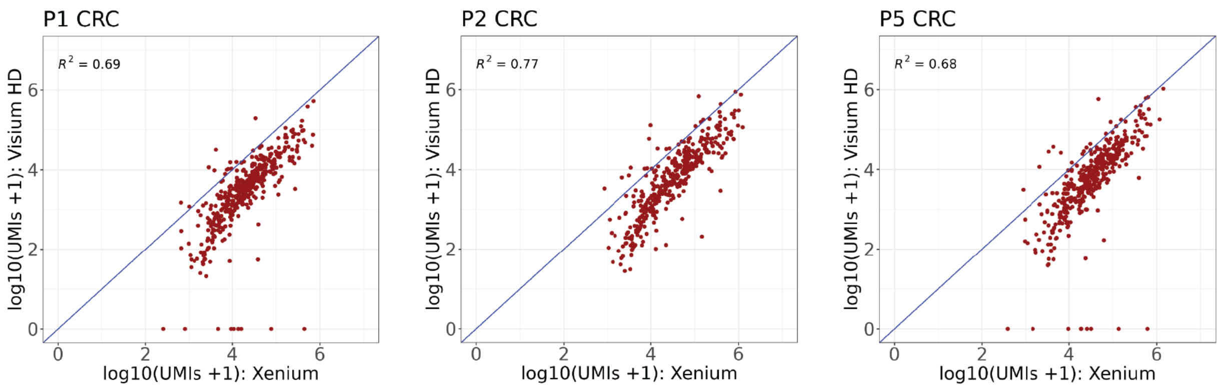


**Supplemental Figure 7. Cross-platforms Sensitivity.** Sensitivity comparisons between Visium HD and Xenium in situ gene expression data have been performed on serial sections from the same colon cancer FFPE blocks in a subset of 3 CRC samples. Plots show per gene pseudo bulk correlation between paired Visium HD (UMI counts) and Xenium in situ (gene counts) data. Xenium is on average 5.7x more sensitive on a per-gene basis than Visium HD for genes included in both panels at the sequencing depth used (range: 1309-1865 reads per 8 µm bin). Sensitivity was calculated by taking the geometric mean of the per gene fold difference between Visium HD and Xenium counts.

We also compared the transcript diversity in the shared region and found that Visium HD exhibited, on average, ~6.5x more transcripts than Xenium.

Total number of transcripts and genes detected in the shared region:

| | Visium HD transcripts | Xenium transcripts | Fold difference in transcripts (HD/Xenium) | Visium HD genes | Xenium genes |
|---|---|---|---|---|---|
| P1 CRC | 107,940,889 | 21,661,383 | 4.98 | 18964 | 422 |
| P2 CRC | 265,645,676 | 41,387,070 | 6.42 | 18072 | 422 |
| P5 CRC | 249,911,348 | 33,040,562 | 7.56 | 18972 | 422 |
| Mean | 207,832,638 | 32,029,672 | 6.49 | 18964 | 422 |

### Consortium Members

**Visium HD Development Team: Consortium of 10x Genomics team members who contributed to the development of the Visium HD technology used in this manuscript.**

Michael Aguilar, Francis Aguisanda, Chaitanya Aluru, Naishitha Anaparthy, Eric Anderson, Melissa Ando, Emil Anthony Gimenez, Joey Arthur, George Auer, Laura Barrera, Alexander Ben, Kheng Boon Tan, Joon Boon Teo, Andrew Broch, Michele Caceres, Matt Cai, Burton Chang, Cynthia Chang, Peggy Chang, Sidharth Chaturvedi, Bhavika Chauhan, Sharon Chen, Yiren Chen, Chris Cheng, Jennifer Cheung, Meii Chung, Nancy Conejo, Julia Cowen, Filip Crnogorac, Eileen Dalin, Heather Danforth, Filip Defoort, Josh Delaney, Xun Ding, Keri Dockter, Cynera Dodati, Zara Doddridge, Sultan Doganay Tuncer, Cassidy Dolstra, Emre Erhan, Michelli Faria de Oliveira, Beevan Gill, Andrew Gottscho, Yang Gu, Zhenping Gua, Yvonne Guan, Anushka Gupta, Lila Haba, Garland Hatten, Alex Hermes, Kendall Hoff, Julia Hsu, Yi Hsu, Tack Huat Lee, Daniel Hung, Jason Hung, Amanda Janesick, Guy Joseph, Govinda Kamath, Sharon Karagozlu, Layla Katiraee, Yarden Killmer Maya, Hanyoup Kim, Sugyeom Kim, Wendy Kim, Kirk Ko, Sreenath Krishnan, Josy Kuriakose, Sural Labha, Ashley Lai, Hank Lai, Shea Lance, Susana Lau Lui, Quynh Le, Jennifer Lew, Dongyao Li, Zhihao Li, Alvin Liang, Fang-Chu Lin, Du Linh Lam, Cheyenne Liu, Guanhong Liu, Peizhi Liu, Ken Lo, Glory Lopez, Han Lu, Zixue Ma, Diego Magdaleno, Areeb Mallick, Liza Man, Sean Marrache, Ray Meury, Glenn McGall, Nabil Mikhaiel, Syrus Mohabbat, Laura Munteanu, Monica Nagendran, Amber Nelson, Muhammad Ng, Michael Noor, Shirlene Ong, Steven Ornellas, Ching Ouh Ng, Juan Pablo Romero, Anuj Patel, David Patterson, Zixuan Peng, Adam Phua, Prislyn Poh, Caio Porto, Deng Qing, Priya Rajendran Vishnu, Matthew Robertson, Guillaume Robichaud, Mike Rose, Janani Sampathkumar, Jerald Sapida, Didem Sarikaya, Michael Schnall-Levin, Paul Seah, Sajan Sebestian, Marco Serra, Preyas Shah, Jyoti Sheldon, Shang Shi, Eric Siegel, Hardeep Singh, Amrit Sinha, Ben Sisserman, Susanne Spielberg, Dhanya Sridharan, Jordan Stefani, Jacob Stern, Arjun Sugumar, David Sukovich, Emily Sun, Dave Tan, Neeraj Tayal, Sarah Taylor, Augusto Tentori, Su Theng Kok, Yen Tren, YJ Tsai, Archit Upadhyay, Miriam Valencia, Neil Weisenfeld, Mia Weng, Dieter Wilk, Stephen Williams, Evan Winget, George Withers, Wyatt Woodson, Zhanyu Wu, Taylor Wyatt, Wanting Xu, Niranjan Yadla, Gokay Yamankurt, Qiao Yan Toh, Yi-Xin Yang, Xin Yao, Ravi Yarlagadda, Nickson Yee, Yifeng Yin, Su Yu, Zahoor Zafrulla, Zhaosheng Zhang, Wei Zhou, Orchid Zhu, Shurong Zhu, Xiaoxuan Zhu.
